# Meta-analysis reveals obesity associated gut microbial alteration patterns and reproducible contributors of functional shift

**DOI:** 10.1101/2022.06.05.494850

**Authors:** Deep Chanda, Debojyoti De

**Affiliations:** Laboratory of Cellular Differentiation & Metabolic Disorder, Department of Biotechnology, National Institute of Technology, Durgapur, West Bengal 713209, India

**Keywords:** Obesity, Gut microbial association, Meta-analysis, Machine learning, Functional contributors

## Abstract

Cohort-specific 16S rRNA sequence-based studies associating gut microbiota with obesity are often marred with contradictory findings regarding community structure and composition leading to “reproducibility crisis” of the signals. Moreover, taxonomic drivers of the obesity-linked gut microbial functional imbalances and their replicability also remains unexplored which should be useful for in-depth understanding of obese host-gut microbiota interaction and, strategizing therapeutics. We addressed these questions through unbiased meta-analysis and further machine-learning validation of 692 curated fecal whole metagenomic sequence datasets from diverse geographical locations. Further, obesity-linked pathway shifts were traced back to their specific drivers by integrating the species and pathway profiles through genomic content of the species. We found reproducible depletion of diversity in obese gut microbiome without any pattern in Firmicutes/Bacteroidetes ratio. Additionally, we also identified obesity-linked robust and reproducible gut microbial species and pathway features. Contributors of these pathway features identified as both dataset-specific and shared across the datasets.

## Introduction

Obesity, addressed as a major global health risk of 21^st^ century has nearly tripled since 1975 (WHO, 2018). Along with genetics (Locke et al., 2015) and epigenetics (Wheeler et al., 2013), a number of other factors like demography (Ogden et al., 2014), appetite regulation (Spiegelman and Flier, 2001), lifestyle (Hankinson et al., 2010; Hu et al., 2003), hormone signalling (Spiegelman and Flier, 2001), etc. play important role in the aetiology of obesity. Recently, independent studies have associated gut microbiome with obesity too (Murphy et al., 2016; Pinart et al., 2022; Turnbaugh et al., 2006, 2008). In alignment to these reports, studies related to fecal microbiota transplantation (FMT) being successful in partially reverting obese associated host metabolic phenotypes, suggests gut microbial alteration is not merely a consequence but rather may be a cause of obesity related metabolic imbalance (Napolitano and Covasa, 2020). In spite of all these remarkable observations, most of the cohort-specific 16S rRNA sequence-based studies associating gut microbiota with obesity are inconsistent in reporting alteration of microbial community structure like diversity, Firmicutes/Bacteroides ratio, etc(Castaner et al., 2018)). These studies also contradict while reporting differentially abundant taxa linked to obesity. Moreover, few efforts to generalise gut microbiome association to obesity by pooling 16s rRNA data from several such studies (Pinart et al., 2022; Sze and Schloss, 2016), fail to provide generalisability at the species and functional level because these were based on low resolution 16s rRNA based classification which had inherent problems with sensitivity and reliability (Durazzi et al., 2021; Poretsky et al., 2014). On the other hand, pooling high-resolution whole genome sequence-based studies with gut microbial species to decipher reproducible pattern associations are absent. Accordingly, the link between the species and pathway shifts they contribute, that may provide in-depth insights into obese host-gut microbiome interactions, remains unexplored too. The knowledge of this connectivity is necessary to pinpoint the species contributors of the functional shifts in order to have a mechanistic understanding as well as to target obesity associated dysbiosis.

To specifically address these issues, in this study, we have curated metagenomic sequences of 692 samples comprising 340 obese and 352 lean control samples from seven different geographical locations, processed, statistically analysed, and subsequently validated. In the process, we initially assessed the species and functional patterns using univariate non-parametric statistical tests for each dataset. Subsequently, we performed random-effect meta-analyses to identify the signature microbial features which were then independently validated through a machine learning analyses. We finally linked the taxonomic and functional profiles which uses a sophisticated genomic content-based pathway copy number information for each species and systematically assessed replicability of the drivers of functional shifts.

Our analyses revealed reproducibly depleted gut microbial diversity in obese individual and no association of obesity with F/B ratio, a previously linked hallmark of obesity. Based on our meta-analysis and machine-learning classification, we could establish a panel of robust, highly associative species as well as pathway signatures. Among the highly predictive robust species biomarkers many of are known to be short-chain fatty acid producers (*Intestinimonas buyriciproducens* and several *Alistipes* sp.) and promoters of gut barrier integrity (*Bifidobacterium longum*). These were further validated independently using a machine learning based classifier. Accordingly, among the several reproducible pathways, we also identified a set of SCFAs biosynthesis pathways to be reproducibly lean-associated. Further, we noticed, shifts in gut microbial functional repertoire are contributed by both dataset-specific and replicable taxa across the datasets. Taken together, we infer that loss of overall microbial diversity and their contributory function, is responsible for obesity associated dysbiosis. Subsequently we identified the taxa and functional pattern linked with lean and obese phenotype. We could also underline the key contributors of functional diversities observed in lean individuals. Our study thus emphasise on how connecting species with the functional shifts is helpful for mechanistic understanding of microbial contribution to obesity as well as lay the foundation for designing therapeutic strategies.

## Methods

### Search strategy and inclusion criteria for public metagenomic sequence dataset selection

Due to unavailability of whole genome sequence (wgs)-based dedicated study on obesity, we considered curation of samples from general cohort-specific studies as well as from case-control studies of other diseases. Thus, we conducted thorough keyword searches in PubMed to retrieve studies with publicly available human fecal shotgun metagenome data from inception till May 2021. We entered the following all-encompassing search terms into PubMed: (“humans”[All Fields] OR “humans”[MeSH Terms] OR “humans”[All Fields] OR “human”[All Fields]) AND (“metagenome”[MeSH Terms] OR “metagenome”[All Fields] OR “metagenomes”[All Fields] OR “metagenomic”[All Fields] OR “metagenomically”[All Fields] OR “metagenomics”[MeSH Terms] OR “metagenomics”[All Fields] OR (“metagenome”[MeSH Terms] OR “metagenome”[All Fields] OR “metagenomes”[All Fields] OR “metagenomic”[All Fields] OR “metagenomically”[All Fields] OR “metagenomics”[MeSH Terms] OR “metagenomics”[All Fields])) AND (“cohort studies”[MeSH Terms] OR (“cohort”[All Fields] AND “studies”[All Fields]) OR “cohort studies”[All Fields] OR (“case control studies”[MeSH Terms] OR (“case control”[All Fields] AND “studies”[All Fields]) OR “case control studies”[All Fields] OR (“case”[All Fields] AND “control”[All Fields] AND “studies”[All Fields]) OR “case control studies”[All Fields]) OR (“computational biology”[MeSH Terms] OR (“computational”[All Fields] AND “biology”[All Fields]) OR “computational biology”[All Fields])) AND (“gastrointestinal microbiome”[MeSH Terms] OR (“gastrointestinal”[All Fields] AND “microbiome”[All Fields]) OR “gastrointestinal microbiome”[All Fields] OR (“gastrointestinal tract”[MeSH Terms] OR (“gastrointestinal”[All Fields] AND “tract”[All Fields]) OR “gastrointestinal tract”[All Fields])) AND (“metabolic syndrome”[MeSH Terms] OR (“metabolic”[All Fields] AND “syndrome”[All Fields]) OR “metabolic syndrome”[All Fields] OR (“colorectal neoplasms”[MeSH Terms] OR (“colorectal”[All Fields] AND “neoplasms”[All Fields]) OR “colorectal neoplasms”[All Fields] OR (“colorectal”[All Fields] AND “cancer”[All Fields]) OR “colorectal cancer”[All Fields]) OR (“atherosclerosis”[MeSH Terms] OR “atherosclerosis”[All Fields] OR “atheroscleroses”[All Fields]) OR “IBD”[All Fields] OR (“arthritis”[MeSH Terms] OR “arthritis”[All Fields] OR “arthritides”[All Fields] OR “polyarthritides”[All Fields]) OR (“obeses”[All Fields] OR “obesity”[MeSH Terms] OR “obesity”[All Fields] OR “obese”[All Fields] OR “obesities”[All Fields] OR “obesity s”[All Fields])).

Additionally, we used CuratedMetagenomicData R package to include more samples in our study (and hence more statistical power) from pre-profiled fecal metagenome sequences and associated metadata (Pasolli et al., 2017). For all the datasets considered in our study, we included samples in our meta-analysis, if the subjects: 1) are adult 2) met the WHO BMI criteria of obese and lean, 3) don’t have any gastrointestinal disease, 4) did not take an antibiotic in preceding three months. In order to reduce biasness due to technical variation, we included fecal shotgun metagenome sequenced with Illumina sequencing platform only. We also ensured that the samples included in our study have important metadata information like “Subject ID,” “SampleID,” “Age,” “Sex,” “BMI,” “Country,” “Sequencing platform,” “Total number of reads,” and “Average read length,”. In case of incomplete metadata, we personally communicated corresponding author(s) to get the missing information. Studies requiring additional ethics committee approvals or authorizations or less than 20 case samples were excluded from our analysis.). The entire schema of study selection and data retrieval is summarized in Figure S1. Samples with 18.5 ≤BMI<25 without any comorbidity information were considered as “control” samples, while BMI greater than 30 was regarded as “obese” samples. The BMI criteria were slightly different for Asia-Pacific populations where samples with 18.5≤BMI<23 and BMI≥25 were considered as “control” and “obese,” respectively (Weir and Jan, 2021).

### Taxonomic and Functional profiling

Initially, taxonomic profiling was performed by MetaPhlAn3 with the “--ignore_eukaryotes” flag to remove contamination (Beghini et al., 2021). Prior to functional profiling by HUMAnN3 (Beghini et al., 2021), *in silico* removal of contaminant host reads were done by aligning the sequences to the human genome (GRCh37/Hg19) by using KneadData integrated Bowtie2 (V.2.3.4.1) tool (Langmead and Salzberg, 2012). Species and functional pathways with relative abundance >0.01% and >0.0001% respectively and prevalence >5% were included in the further analyses.

### Statistical Analysis

All the statistical analyses were carried out within R 4.0 (https://www.R-project.org/) unless otherwise stated. We identified significantly differentially abundant Taxonomic and functional features between the two groups in each dataset using linear discriminant analysis effect size (LEfSe) (https://huttenhower.sph.harvard.edu/lefse) (Segata et al., 2011). LEfSe uses a non-parametric Kruskal-Wallis (KW) sum-rank test to identify significantly different features along with their respective effect size (LDA score). LDA score ≥2.0 and ≤ -2.0 with a *p-value* <0.05 was considered significant. We also performed multivariable analyses in presence of potential confounders. For the multivariable analyses, we used Microbiome Multivariable Association with Linear Models (MaAsLin2) R package to fit general linear models on our data (Mallick et al., 2021). MaAsLin2 was run first, in the absence of any covariate to compute “crude coefficients” and then in the presence of covariates (age and sex) to compute “adjusted coefficients”. Consequently, we used a linear regression model to analyze the relationship between MaAsLin-derived crude and age-, sex-adjusted coefficients to assess if the crude coefficients are meaningfully affected by the potential covariates. For the meta-analysis, we first, converted the feature relative abundances into arcsine-square root-transformed proportions and used *escalc* function from the *metafor* R package (Viechtbauer, 2010) to compute standardized mean difference (SMD) in terms of Cohen’s D. Consequently, random-effects model estimates were calculated from SMD using rma.mv function of the same package. Finally, meta-analyses results were visualised with the help of *ggforestplot* R package. P-values obtained from random effects were adjusted using the FDR method to adjust for multiple hypothesis testing. Any species present in all datasets with an effect size greater than 0.2 or less than -0.2 and FDR <0.1 was considered reproducible and significantly differentially abundant taxa across the datasets. Similarly, for function level meta-analysis, a function was considered reproducibly significant when present in all datasets with an effect size greater than 0.35 or less than -0.35 at an FDR <0.05. P value <0.05 was considered significant for all the statistical analysis reported here. The percentage of variation across studies for all the meta-analyses reported here was determined using *I*^2^ statistics as well as Cochran’s Q-test for estimating statistical significance of heterogeneity (<u>Higgins et</u> <u>al.,2022)</u>. We considered *I*^2^<60% as acceptable heterogeneity in our random-effect meta-analyses.

### Taxonomic and Functional diversity and Firmicutes/Bacteroidetes Ratio Analysis

Alpha-, beta-diversity, gene richness and Firmicutes/Bacteroidetes (F/B) ratio analyses were performed for each dataset. Prior to diversity analyses, for the sake of normalization, all the samples were rarefied to 90% of the lowest sample depth in a dataset specific way. Rarefaction and diversity analyses were performed by phyloseq 1.26.1 R package (McMurdie and Holmes, 2013). We calculated three standard metrics of alpha diversity: Shannon’s and Simpson’s diversity indices for species, and observed richness for gene family. Beta diversity was estimated through Bray-Curtis dissimilarity metric. Statistical significance of the difference in alpha and beta diversity was tested independently between the two groups by the Wilcoxon Rank Sum test. Next, we summarised the taxonomic profiles of the samples up to the phylum level and calculated the ratio of the relative abundances of Firmicutes and Bacteroidetes (F/B ratio). Wilcoxon Rank Sum test was used to estimate the statistical significance of F/B ratio between the groups. We also performed the same statistical analysis for assessing significance of difference in gene richness between the control and obese groups.

### Random forest-based classifier

The machine learning analyses were performed using the MetaML tool that used taxonomic and functional relative abundances as input. Random forest (RF) classifier was implemented for each dataset using MetaML machine learning tool (Pasolli et al., 2016). We performed three different modes of prediction, where we tested inside-dataset prediction capability (Cross Validation, CV), across-dataset prediction performance (Cross Study Validation, CSV), and Leave One Dataset Out (LODO) approach that mimics the meta-analysis. We assessed inside-dataset prediction accuracies by 10-fold cross validation where a balanced proportion of the obese and control samples were used from same dataset for both learning and validation in each fold. Each 10-fold cross-validation was iterated 20 times, and the average accuracy from the 200 validation folds was reported. Transportability of the model across datasets to evaluate the generalisability and to find whether the taxonomic and functional features are predictable across the datasets were accessed by performing Cross Study Validation (CSV). In CSV, the model was trained on a dataset and validated on a completely independent dataset to evaluate the reproducibility. CSV was repeated for all combinations of independent datasets. Moreover, we also accessed the predictability of the model when trained on multiple datasets using Leave One Dataset Out (LODO) approach also known as hold-out study. In this setting, the model was learned on pooled samples from all the datasets except the one that was used for model validation. The test and the validation dataset for the purpose was chosen independently. Furthermore, the specificity of the prediction was assessed by randomly shuffling the class labels of the samples and comparing the result with the original class. We used AUC value as metric for the prediction performance. The potential discriminative features for classification were further obtained through internal feature ranking from LODO study. Briefly, we first, computed the minimum number of features required for the classification by sequential adding of features and subsequently estimating the AUC values. We ranked the minimum discriminative features capable of distinctively classifying the samples for each LODO analysis. Subsequently, we computed the average rank for these features.

### Identification of reproducible contributors of signature pathway shifts

To pinpoint the taxonomic contributors of the functional shifts, we used the FishTaco tool that systematically integrates taxonomic and functional profiles to trace back to specific contributor taxa according to the protocol described in (Manor and Borenstein, 2017) with minor modifications. Briefly, we first manually curated all the genes and their respective copy numbers for all the species involved in our study from Integrated Microbial Genomes and Microbiomes (IMG) database version 5.0 (https://img.jgi.doe.gov/). We subsequently constructed the gene to functional pathway mapping file using IMG and MetaCyc database version 23.1 (https://metacyc.org/) depending upon the presence-absence pattern of a gene in a pathway. Pathway copy number was computed from the gene to pathway mapping files and the average Gene copy number of the genes constituting the pathway. Finally, FishTaco was run separately for each independent dataset to quantitate the contribution of individual taxa to individual functional shifts. Finally, contributors of 18 functional shifts as obtained previously from meta-analysis were retrieved.

## Results

### Search result and sequence dataset selection

Pooling data from independent studies improve statistical power due to greater sample size and thus can find underlying signal which otherwise would remain unnoticed (Cohn and Decker, 2003)(Jackson and Turner, 2017). Hence, we tried to include as many samples as possible representing different ethnic groups and performed further analyses on the aggregated whole metagenomic datasets as detailed in the analysis workflow (Figure 1). The search strategy as described in the method yielded 200 publications on human gut microbiome related studies. A further screening of title and abstract, short-listed 92 publications for their potential inclusion in our study. Based on the inclusion criteria as mentioned in method section, a detailed evaluation of the studies, was performed and we finally included 6 studies in our meta-analysis which accounted for a total 404 samples including 196 obese and 208 control samples. These studies include samples from 6 different countries (China, Denmark, Ireland, Japan, Kazakhstan and Sweden) (Figure S1). Additionally, we curated pre-profiled samples that fulfilled inclusion criteria from a Great Britain (GBR: 144 obese and 144 control samples) cohort using CuratedMetagenomicData R package. Thus, taken together, we included a total of 692 samples which included 340 obese and 352 control samples from 7 different countries (Table 1 and Figure S1). For our meta-analysis, we downloaded approximately 4TB fastq data and related metadata from Sequence Read Archive (SRA) from the links provided in the original article (except for the GBR dataset). General characteristics of the datasets and associated metadata are summarised in Table 1 and Table S1 respectively.

**Figure 1.**
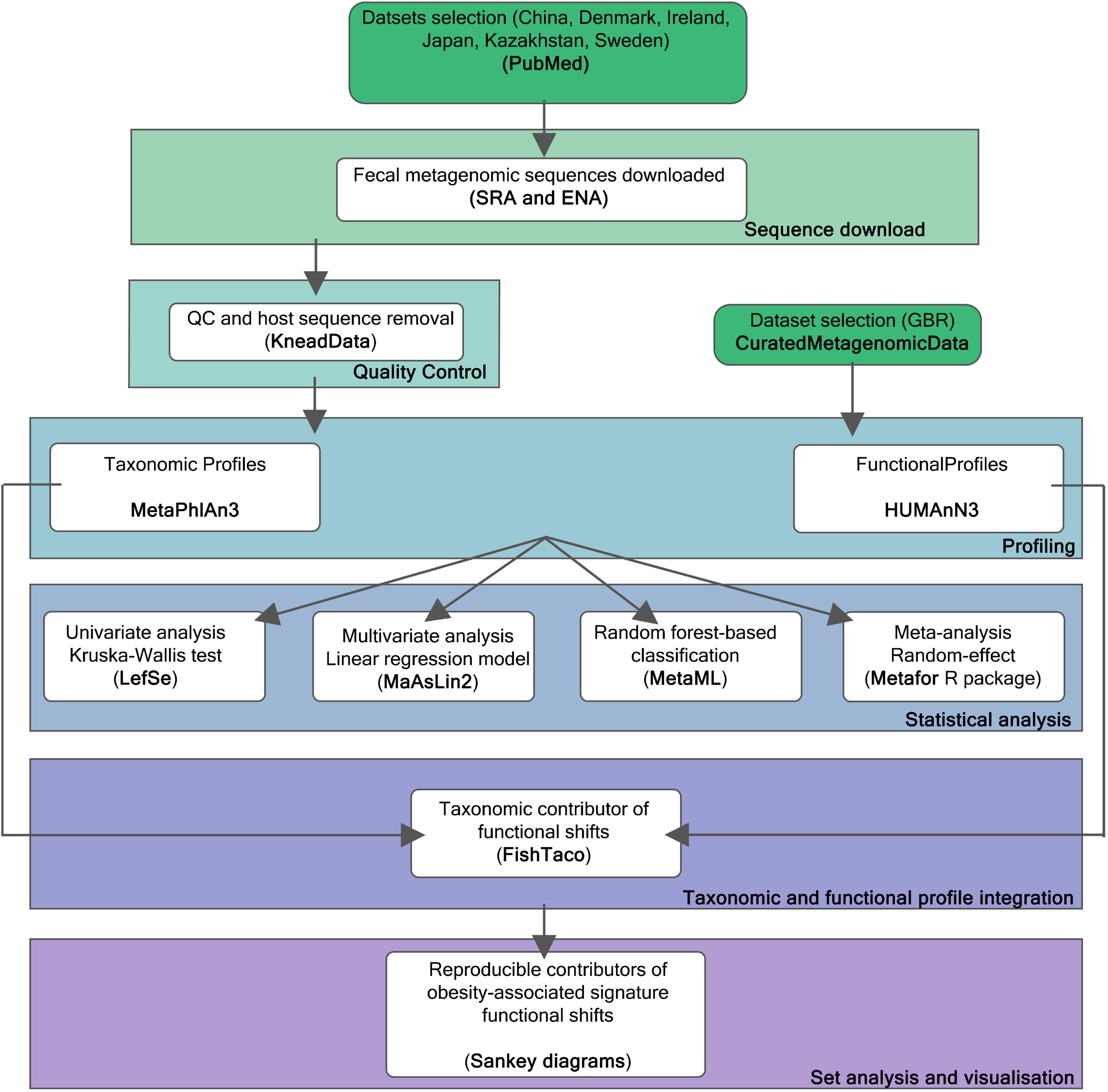
Analysis Workflow. Overall flow chart of our analysis for curating, profiling and analysing lean and obese fecal metagenomic fastq sequencing data.

**Table 1:** General characteristics of the datasets used in this study.

| NCBI BioProject ID | Groups (n) | Age (average ± s.d.) | BMI (average ± s.d.) | Female/male | Country | No. of reads x (10<sup>9</sup>) |
| --- | --- | --- | --- | --- | --- | --- |
| PRJEB1786<br>(Karlsson et al., 2013) | Control (16)<br>Obese (20) | 70.5 ± 0.72 | 28.9 ± 6.69 | 36/0 | Sweden | 1.06 |
| PRJDB4176<br>(Yachida et al., 2019) | Control (46)<br>Obese (45) | 62.9 ± 11.82 | 24.4 ± 3.46 | 37/54 | Japan | 4.45 |
| PRJEB2054<br>(Qin et al., 2010) | Control (37)<br>Obese (31) | 56.1 ± 10.32 | 27.4 ± 6.07 | 35/33 | Denmark | 4.53 |
| PRJEB17632<br>(Kushugulova et al., 2018) | Control (25)<br>Obese (23) | 48.7 ± 10.99 | 27.4 ± 4.89 | 32/16 | Kazakhstan | 2.57 |
| PRJEB21528 | Control (45) | 61.1 ± 9.5 | 25.9 ± 9.04 | 53/30 | China | 4.53 |
| (Jie et al., 2017) | Obese (38) |  |  |  |  |  |
| PRJEB37107<br>(Ghosh et al., 2020) | Control (39)<br>Obese (39) | 76.4 ± 7.98 | 28.2 ± 6.73 | 43/35 | Ireland | 3.32 |
| PRJEB39223<br>(Pasolli et al., 2017) | Control (144)<br>Obese (144) | 45.2 ± 11.87 | 27.5 ± 8.20 | 231/57 | Great<br>Britain<br>(GBR) | 8.52 |
| Total | Control (352)<br>Obese (340) |  |  |  |  | 29 |

### Altered species and gene diversity is associated with obese gut with no significant change in F/B ratio

Substantial discrepancies with respect to obese associated gut microbial species diversity, gene richness and F/B ratio were reported in many previous cohort-specific studies (Castaner et al., 2018; Magne et al., 2020). In order to assess whether a pan-ethnic reproducible pattern exists, we initially analysed these parameters in individual datasets using univariate analysis. From univariate non-parametric test on Shannon and Simpson alpha diversity metrics, we found a consistent reduction in gut microbial diversity in obese individuals except the Chinese dataset (Figures S2A and S2B). In alignment to this, we could observe that the median alpha diversity indices decreased in obese subjects in all datasets except in Chinese. Although this decrease was consistent among datasets it was not statistically significant except for Kazakh (Shannon: p=0.045) and GBR datasets (Shannon, p=0.021; Simpson, p=0.0061) (Table S2). In alignment to the above observation, we also found the obese individuals have reduced gene count than the lean controls except the Irish dataset among which this reduction is significant in three datasets (GBR, Japan and Sweden) (Figure S2C and Table S2). Furthermore, we could not find any consistent pattern in the F/B ratio and also their alteration across the datasets were insignificant (Figure S2D and Table S2). We were also interested to see how the gut microbial composition differs between the individuals of a group. For each dataset, first, we calculated compositional dissimilarity (beta diversity) within the control group (“intra-control” beta diversity) and within the obese group (“intra-obese” beta diversity). We subsequently performed Wilcoxon rank sum test to access the difference of the beta diversity between the groups. It was observed that the intra-control beta diversity was higher in majority of the datasets and was significant in Ireland, Japan and Kazakhstan datasets (p<0.05) (Figure S2E and Table S2). As a next step, we calculated beta diversity between the individuals of obese and control group as “inter-group” diversity and compared them with “intra-control” diversity as well as with “intra-obese” diversity obtained earlier. Expectedly, inter-group beta diversity was found to be higher than both intra-control and intra-obese beta diversity in majority of the datasets although was not significant in all (Figure S2E).

In all these univariate statistical analyses, despite some significant alterations, we observed a lot of inconsistencies across datasets and therefore we could not deduce any reproducible pattern. Thus, we choose to perform random-effect meta-analysis which pools data across the datasets to increase number of observations. Thus, meta-analyses are reported to have higher statistical power and accuracy to find underlying pattern across datasets (Cohn and Decker, 2003). Pooled analysis on the standardized mean difference (Cohen’s d) of the alpha diversity indices confirmed its significant reduction across the datasets [ For Shannon, meta-analysis coefficient estimate (μ) =0.22, 95% confidence interval (0.06, 0.36), p=0.004) and For Simpson (meta-analysis coefficient estimate (μ) =0.19, 95% confidence interval (0.04, 0.34), p=0.012)] with no significant heterogeneity (Shannon: *I*^2^=0%, p=0.755, Q-test; Simpson: *I*^2^=0%, p=0.841) (Figure 2A and Table S3). In parallel to this, meta-analysis with gene richness data revealed significant reduction with negligible heterogeneity (*I*^2^=4.28%; p=0.52, Q-test; meta-analysis coefficient estimate (μ) =0.19, 95% confidence interval (0.03, 0.34), p=0.01) across the datasets (Figure 2A). Similar to univariate analysis, our poled analysis of the F/B ratio was represented with a small effect size, failed to retrieve any reproducible pattern associated to phenotype (Figure 2A and Table S3). In alignment to our finding, there are reports questioning the validity of using the F/B ratio as a hallmark of obesity (Magne et al., 2020; Sze and Schloss, 2016). We further performed random effect meta-analysis to assess if a pattern exists with respect to the beta diversity alteration across the datasets. Again, we did not find any reproducible association when intra-control and intra-obese beta diversity indices were compared through meta-analysis. We also probed the reproducibility of differences of inter-group beta diversity with intra-control and intra-obese beta diversity by random-effect meta-analysis (Figure 2B). We observed that in both of these, the difference was insignificant (Table S3) with the inter-group beta diversity being slightly higher than intra-obese beta diversity. This observation can be partly explained by higher compositional variation in gut microbial community within lean subjects with respect to the obese individuals (Figure 2B). Furthermore, in order to find if our meta-analysis is confounded by the potential covariates like age and sex, we performed multivariate analysis on a per dataset basis with MaAsLin2. When the crude- and adjusted (age and sex) coefficients were fit on a linear model, we observe the covariates did not meaningfully affect the crude coefficients (Figures S3A-S3C). It was also confirmed by performing meta-analysis using crude- and adjusted coefficients obtained across all datasets (Figures S3D and S3E).

**Figure 2.**
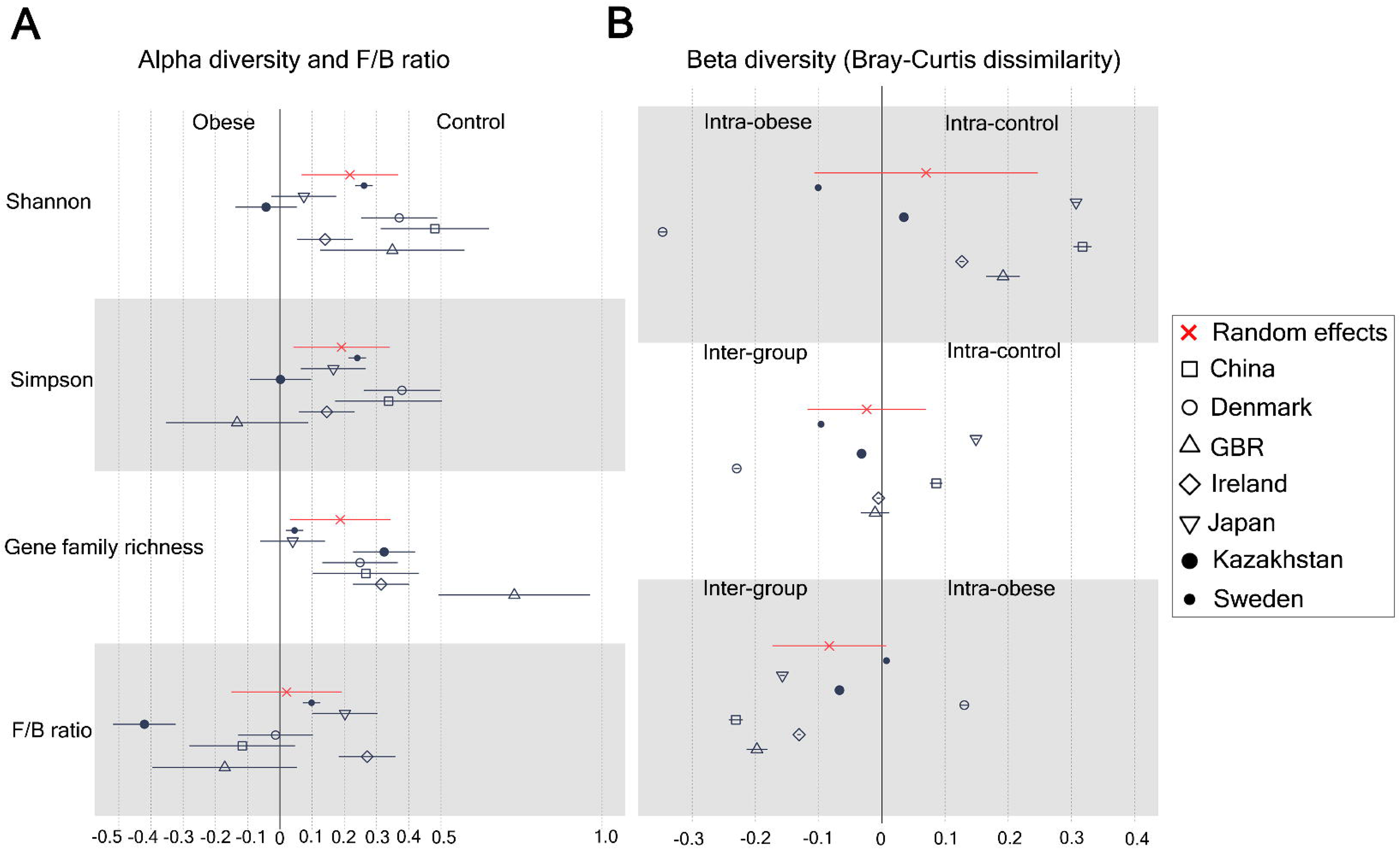
Meta-analysis of Common Metrics Indicating Taxonomic and Functional Structure of the Community. (A) Measures of alpha diversity at taxonomic (Shannon, Simpson) and functional (gene richness) level and Firmicutes/ Bacteroidetes ratio. (B) Beta diversity metrics (Bray-Curtis dissimilarity) among and between control and obese groups.

### Identification of Obesity-linked signature taxa

To identify obesity-linked differentially abundant taxa across the datasets, initially, we performed a non-parametric Kruskal-Wallis (KW) test by LEfSe on the species relative abundance profiles between the two groups considering the non-normal distribution of the microbiome composition (Figures S4A-S4G). We observed that majority of the species are dataset specific with 3 lean enriched species (like *Anaeromassilibacillus* sp. An250, *Bacteroides nordii, Intestinimonas butyriciproducens)* being present in 3 out of 7 datasets (Figure 3A). This observation is particularly important as we could find extensive reports of associating them negatively with BMI and protective roles against metabolic syndrome, implying that they could be markers of a healthy gut (Brahe et al., 2015; Tagliamonte et al., 2021; Treichel et al., 2019). Taken together, our univariate analysis failed to identify any reproducible taxa across the datasets as no species were found to be differentially abundant in at least 4 out of 7 datasets (Figure 3A). The absence of reproducibility may be due to variations in factors like diet, ethnicity, biogeography, host genetics, sample size, and sample processing that impart biases in individual datasets.

**Figure 3.**
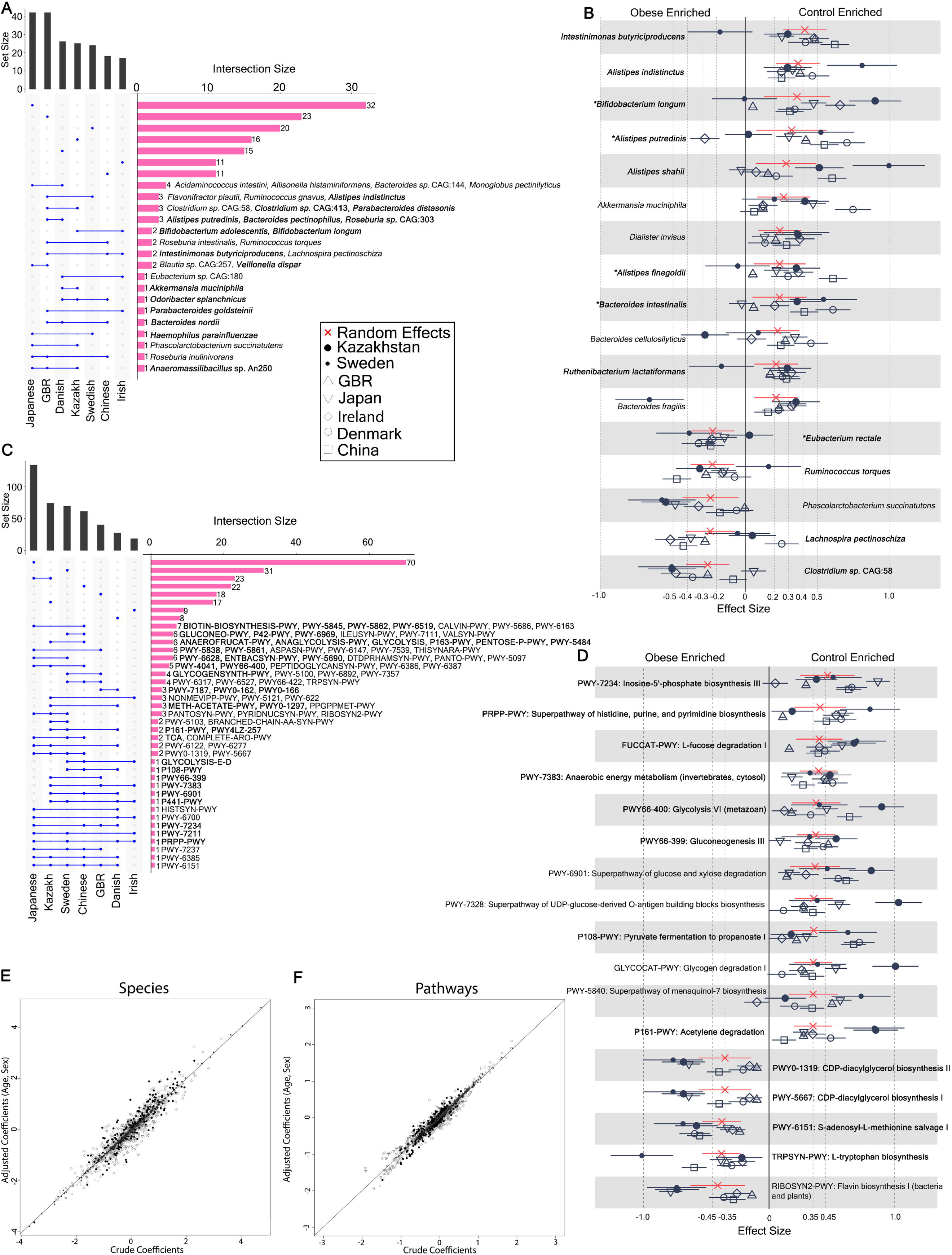
Reproducible Taxonomic and Functional Microbial Biomarkers Across Datasets When Comparing Obese to Lean Controls. (A) UpSet plot showing number of taxonomic biomarkers identified using LEfSe on MetaPhlAn3 generated species profiles shared among the datasets. Species marked in bold are control-enriched. (B) Pooled effect sizes for the 17□significant species with FDR cut-off less than 0.1. Red lines represent the 95% confidence interval for the random effects model estimate. Species marked in bold refer to the biomarkers which were also validated from machine learning analysis (Figure 5). Species marked in ‘*’ denote reproducible contributors of the obesity-linked gut microbial signature pathways as identified from FishTaco analyses (Figure 6). (C) UpSet plot showing number of gut microbial pathway biomarkers identified using LEfSe on HUMANn3 generated pathway profiles shared among the datasets. Pathways marked with bold letters are control-enriched. (D) Pooled effect sizes for the 17□significant pathways with FDR cut-off less than 0.05. Red lines represent the 95% confidence interval for the random effects model estimate. Pathways marked in bold were confirmed by independent machine-learning analysis (E) (E-F) Scatter plot of crude and age, sex adjusted coefficients obtained from linear models using (E) MetaPhlAn3 species abundances, and (F)HUMANn3 pathway abundances.

To explore the reproducibility further, we pooled species level relative abundance data by random effect meta-analysis which identified 23 species to be replicable across all the datasets at FDR <0.1 (Table S4.1). We identified 7 bacterial species when used an effect size cut off of ±0.25 at FDR <0.1 (Figure 3B). Out of which 6 were control enriched: *Intestinimonas butyriciproducens* (μ=0.41, 95% CI=(0.26, 0.56), FDR =1.10E-05)*, Alistipes indistinctus* (μ=0.36, 95% CI=(0.21, 0.52), FDR =0.0001)*, Bifidobacterium longum* (μ=0.36, 95% CI=(0.13, 0.59), FDR =0.050)*, Alistipes putredinis* (μ=0.32, 95% CI=(0.08, 0.56), FDR=0.068)*, Alistipes shahii* (μ=0.28, 95% CI=(0.07, 0.49), FDR =0.068)*, Akkermansia muciniphila* (μ=0.27, 95% CI=(0.08, 0.45), FDR =0.055) and 1 obese enriched: *Clostridium sp* CAG:58 (μ=-0.26, 95% CI=(−0.11, -0.41), FDR =0.026). We did not find significant heterogeneity across the datasets (Q test, p-value>0.05, *I*^2^<60%) (Table S4). Earlier studies reported that some of the identified lean enriched species, are the ones that also became prevalent when obese individuals took Mediterranean diet, suggesting their protective role (Tagliamonte et al., 2021).

Altogether, at FDR < 0.1 and effect size cut off ±0.2, we identified a set of 17 species reproducible across the datasets majority of which are control-enriched and previously associated with healthy metabolic status and BMI (Figure 3B and Table S4.1) (Liu et al., 2017; Testerman et al., 2021; Verdam et al., 2013; Zhernakova et al., 2016). On the other hand , the obese-enriched taxa were also linked to impaired host energy metabolism or high BMI (Liu et al., 2017).

### Identification of Obesity-associated reproducible pathways

Although univariate non-parametric test failed to detect reproducibility at the taxa level, LEfSe analysis on pathway level (Figure S5) revealed eight gut microbial metabolic pathways are significantly differentially abundant in at least four datasets (Figure 3C). Among them, the *S*-adenosyl-L-methionine salvage I pathway (PWY-6151) was significant in six datasets except the Irish dataset while, CDP-diacylglycerol biosynthesis I (PWY-5667), peptidoglycan biosynthesis III (mycobacteria) (PWY-6385), *myo*-, *chiro*- and *scyllo*-inositol degradation (PWY-7237), CDP-diacylglycerol biosynthesis II (PWY0-1319), superpathway of *N*-acetylneuraminate degradation (P441-PWY), superpathway of histidine, purine, and pyrimidine biosynthesis (PRPP-PWY), inosine-5’-phosphate biosynthesis III (PWY-7234) are each significant in four datasets. Rest are either shared by fewer datasets or are exclusively dataset-specific. The result reasonably indicates that unlike taxa, some level of reproducibility exists at the pathway level, which directed us to further explore this through random-effect meta-analysis as it has better statistical power.

From meta-analysis, we identified 70 pathways enriched or depleted in obesity at FDR<0.05 (Table S4.2). Among these, we focused on a list of 17 pathways which were retrieved with stringent criteria (FDR<0.05; Effect Size cut off ±0.35) (Figure 3D, Table S4.3). Of which, 12 pathways were control enriched, and only five were obese enriched. Notably, control enriched pathways are involved in purine and pyrimidine biosynthesis, fermentation, carbohydrate metabolism, nucleotide sugar biosynthesis, Vitamin *K*_2_ (menaquinol) biosynthesis (Table S4.3). Among the control enriched reproducible pathways purine and pyrimidine biosynthesis pathways show highest effect sizes. These pathways were also identified as significantly control-enriched in majority of the datasets (Figure 3C). Our finding supports earlier studies reporting negative association of BMI with purine-, pyrimidine nucleotide biosynthesis, glycolysis, gluconeogenesis, menaquinone biosynthesis (Liu et al., 2017). Furthermore, the obese enriched pathways are involved in cofactor biosynthesis, amino acid biosynthesis, phospholipid biosynthesis. S-adenosyl-L-methionine cycle I pathway shows highest effect size. Per dataset univariate analysis revealed this pathway significantly enriches in obese group in 6 out of 7 datasets. Among the other obese enriched reproducible pathways CDP-diacylglycerol biosynthesis pathways I and II were also found to be significantly enriched in majority of the datasets as identified from univariate non-parametric analyses. Thus although, from the univariate non-parametric analyses we could not find any signature pattern at the taxa and functional level random effect meta-analyses proved effective in identifying underlying patterns.

Considering the limitations of assessing the effect of confounders in the univariate analysis (both and LEfSe and meta-analysis), we performed multivariate analysis with both taxa and functional features by MaAsLin2. First, for each dataset, we calculated crude- and adjusted coefficients respectively in the absence and presence of the potential covariates (age, sex). When the two coefficients were fit on a linear model, we observed the covariates did not meaningfully affect the crude coefficients of meta-analysis (Figures S6A-S6D). Additionally, to access whether signature features are confounded by ethnicity, we evaluated relative abundance profiles of the signature features through MaAsLin2 to find the crude and ethnicity-adjusted coefficients. Subsequently, when these coefficients were fit on a linear model we noticed negligible influence of the ethnicity on crude coefficients (Figures S6E and S6F). This observation strengthens cross-dataset reliability of the signature features we deciphered.

### A machine-learning based classifier validates taxa and functional pattern

To check the validity of the pattern obtained through meta-analysis, we resorted to explore the pattern independently using a machine-learning based approach, which are also very powerful to decipher patterns. We set out to validate if the signature of independent datasets is transferable to other studies and if pooling data across studies is more efficient to predict an underlying pattern. Thus, we performed three sets of prediction tasks; Cross Validation (CV), Cross Study Validation (CSV), Leave One Dataset Out (LODO) to check which is the best performing model (see Materials and Methods). For the purpose, we implemented random forest (RF) as backend classifier through MetAML machine learning package as it outperforms other machine learning algorithms with microbiome data (Pasolli et al., 2016). First, we performed cross validation (CV) experiment, by training and validating the RF classifier with species relative abundance data on the same dataset and found the prediction performance ranges between AUC (area under the receiver operating characteristic curve) value 0.57 and 0.73 with median value 0.61 (s.d. ± 0.07) (Figure 4A). We observed training and validation on GBR dataset results in highest predictive accuracy (AUC = 0.73) probably because the dataset has highest number of samples (N = 288) along with high quality of the data. Sufficient accuracy level was also achieved when CV was performed on Danish (AUC = 0.72) and Chinese (AUC = 0.67) datasets. However, despite a comparable sample size (N = 78), CV in Irish dataset resulted in lowest AUC (0.57). We observed AUC values varying over a wide range between 0.38 and 0.72 (Figure 4A). In all except Irish dataset, the intra-dataset prediction accuracy found to be higher than the median accuracy generated from the CSV specific to that dataset. However, the dataset-specific median accuracy obtained from CSV ranged between 0.47 and 0.66 (median AUC = 0.58; s.d. ± 0.06) overall implying a reduced accuracy in CSV in comparison to CV. Since, dataset from different ethnicity had their respective intricacies in features, prediction ability was low when the “training” was done on single dataset followed by validation in the same dataset (CV) with respect to the validation on different datasets (CSV), possibly due to under fitting.

**Figure 4.**
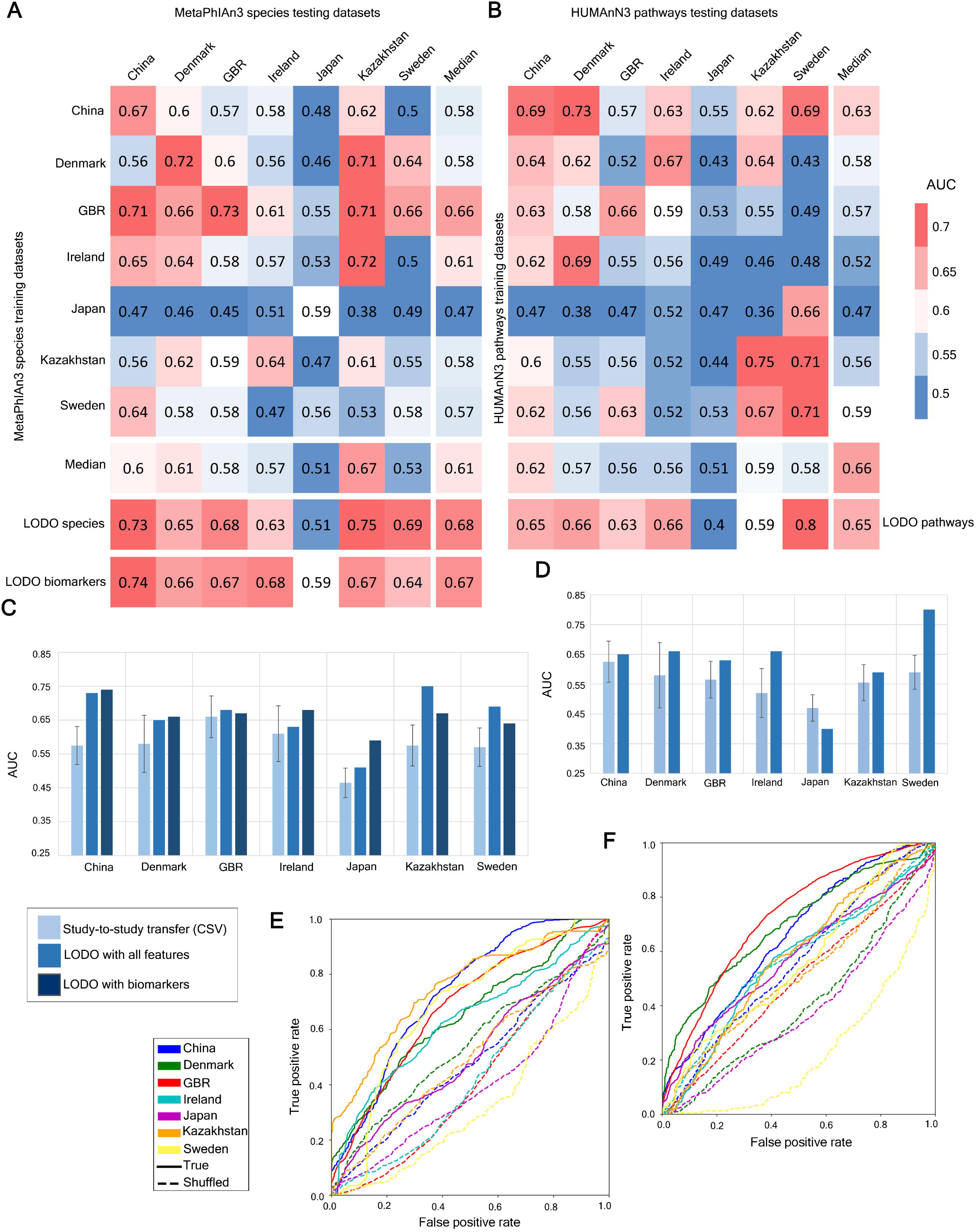
Assessment of prediction performances of the gut microbiome for class prediction within and across cohorts. (A-B) Matrix depicting prediction performances in terms of AUC values obtained from MetaML using random-forest as backend classifier on (A) species-level relative abundances and (B) pathway-level relative abundances. Diagonal values refer to average AUC obtained from 10 fold cross validations iterated 20 times. Off-diagonal values report AUC values generated by training classifier on a dataset present in a row and validated on a dataset in the corresponding column. LODO rows refer to the accuracy achieved by training the classifier on all but the dataset of the corresponding column with relative abundances of species (LODO species), pathways (LODO pathways) and biomarkers (LODO biomarkers). (C-D) Comparison of accuracy achieved when classifier is trained on species (C) and pathway (D) relative abundances from multiple datasets (LODO) with respect to the classifier trained in a single dataset and transported to others datasets (CSV). (E-F) Average of ROC curves obtained (over fold) for LODO analyses in the presence of true and shuffled class levels showing specificity of the prediction by the classifier with species (E) and (F) pathway profiles. Consistently higher accuracy is achieved with true class levels as compared to the shuffled ones for all datasets.

We also performed similar machine-learning based predictions (CV and CSV) with functional pathway features. We found AUC values in CV ranged between 0.47 and 0.75 with a median AUC value 0.66 (s.d. ± 0.1). Except Ireland (AUC = 0.56), Japan (AUC = 0.47) and Denmark (AUC = 0.62) all achieved sufficient classification accuracy (Figure 4B). Among them, CV with Kazakh (AUC = 0.75) and Japanese (AUC = 0.47) datasets achieved the highest and lowest prediction accuracies respectively. Besides, when classifier was trained on a dataset and validated on others, we observed a reduced median accuracy than when it was validated on itself (CV) for all datasets (except Japan). However, the dataset-specific median accuracy obtained from CSV ranged between 0.47 and 0.63 (median AUC = 0.57; s.d. ± 0.05) overall implying a reduced accuracy in CSV in comparison to CV as also found with species features.

Altogether, AUC values are comparatively higher in CV than CSV, suggesting that intra-dataset prediction-validation is specific to dataset and microbial signature are poorly transferable across the datasets. This is line with our dataset specific univariate non-parametric statistical analyses where significantly differentially abundant species in one dataset were rarely found to be reproducible on other datasets.

### Case versus control discrimination was improved across datasets in LODO setting

In order to access the validity and reproducibility of our meta-analysis result, we mimicked the meta-analysis by pooling all but one dataset for training, and used the same for validation to evaluate generalisability of the model (LODO). Thus, 7 sets of LODO experiments, each for one dataset was performed for species and pathway features separately. When LODO was performed with species abundance, expectedly, in line with our meta-analysis, we observed a substantially improved accuracy in transferability of the learning model than that of the CSV setting for all datasets (Figure 4A and 4C). LODO classification on Kazakh and Chinese datasets achieved the best two prediction accuracy (AUC = 0.75 and 0.73 respectively) (Figures 4A and 4C). Substantial level of discriminatory power was also observed for other three datasets: Sweden (AUC = 0.69), GBR (AUC = 0.68) and Denmark (AUC = 0.65). However, in LODO, we did not observe considerable discriminative performance in Ireland (AUC = 0.63) and Japan (AUC = 0.51), which were still better than the average prediction accuracy score achieved in their respective median CSV (Figure 4C). Additionally, we observed median accuracy of LODO (AUC = 0.68) settings exceeded the median accuracy obtained from CV (AUC = 0.61) experiments. Overall, inclusion of more datasets in training increases host population diversity which thereby improves generalizability of classifiers as we evidently found the exercise at the species level substantially and consistently outperforms CV and CSV.

To evaluate how inclusion of population diversity during the training phase influence the overall prediction accuracy, we computed the AUC values for each dataset (validation phase) with while gradually increasing number of datasets one at a time (Figure S7 A-S7G). We noticed a sharp rise (5% for GBR dataset to 21% for Swedish dataset, Figure S7H) in the median AUC value while including from one to two training datasets for majority of the validation datasets (China, Denmark, GBR, Kazakhstan, Sweden) in LODO. Further inclusion of datasets resulted into less prominent changes in median AUC values. Therefore, addition of large, heterogeneous training datasets promotes efficient and accurate discrimination between the classes.

Prompted by the success at the taxa classification we also performed LODO exercise involving the pathways. With the pathways too, the AUC values improved with respect to the CSV analyses in all dataset except for Japan (Figure 4B and 4D). Highest accuracy was observed with Sweden dataset (AUC = 0.8) whereas, followed by China (AUC = 0.65), Denmark (AUC = 0.66) and Ireland (AUC = 0.66) datasets, where sufficient discrimination between the classes were also observed (Figures 4B). However, unlike species level, we found a slight reduction of median accuracy of LODO analyses (AUC = 0.65) than that obtained from CV experiments (AUC = 0.66). Although we observed pooling pathway abundance data by meta-analysis captures the signature more effectively than dataset-specific non-parametric analysis, the same was not true in machine learning experiments. Thus, for the pathway level analysis, different machine learning algorithm or even deep learning may be useful. Still, the AUC values obtained by using same classifier with shuffled class labels found to be significantly lower than those obtained from original class labels with both taxa and functional features in LODO analysis, highlighting the specificity of the machine learning model for class discrimination (Figures 4E and 4F and Table S5). This observation implies substantial level of association of specific microbial features with obesity.

### Validation of obesity associated signature species using feature ranking

As LODO mimicking meta-analysis outperforms CSV with both species and pathway relative abundance data, we used it to validate obesity-associated taxa from our meta-analysis (Figures 3B and 3D). For the purpose, we looked for a minimum set of features that can attain an AUC comparable to that of using complete feature set. While calculating the AUC values with increasing number of features, we observed a marginal increase in average AUC (1.5% to 4.5%) in majority of the datasets when all species were considered for classification instead of using as few as 32 species (Figure 5A). Similarly, for the pathway level, again only a slight improvement (0% to 3%) in average AUC values were observed in all except Sweden dataset (12%) (Figure 5B). It suggests that a finite number of species or pathway features can explain obesity-associated pattern and thus meta-analysis results can be validated by comparing with highly predictive features obtained from LODO analyses.

**Figure 5.**
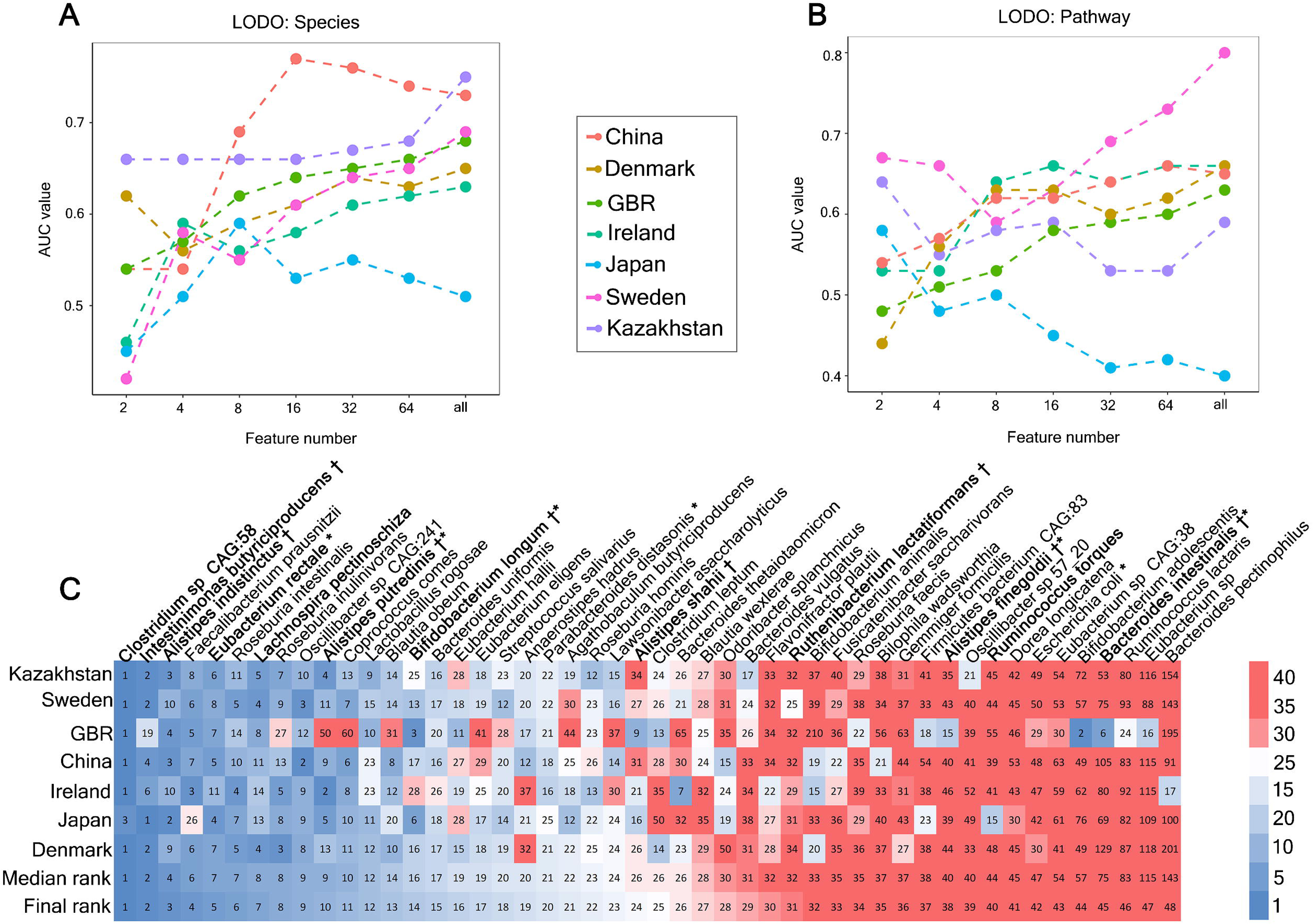
Identification of minimal features required for class prediction and their dataset-specific ranking. (A-B)Predictive accuracy with increasing number of (A) species (B) pathway features obtained by retraining the classifier on the top-ranked features identified through first random forest classifier in LODO analyses. (C)Representation of the rank matrix along with median and final rank of top 32 species in each LODO validation. Species marked in bold denote biomarkers as also obtained from species-level meta-analysis. ‘†’ marked species are the control-enriched biomarkers. ‘*’ indicates species which were obtained as reproducible contributors of obesity-linked signature pathway shifts from FishTaco (Figure 3 and Figure 6).

In order to validate our meta-analysis result, first, we took 32 top-ranking species from each LODO run and averaged their ranking and looked for if they are comparable to our meta-analysis result. Expectedly, most of the species obtained from meta-analysis were also found in the list of top features required for discrimination of the two classes (Figures 3B and 5C and Table S4.1). As expected, *Clostridium* sp. CAG:58 which is an obese-enriched species with highest effect size also shown best discriminatory ability with highest average rank. Moreover, two control-enriched species *Intestinimonas butyriciprioducens* and *Alistipes indistinctus* with highest effect sizes stood second and third highest respectively. Besides, *Eubacterium rectale, Lachnospira pectinoschiza, Alistipes putredinis, Bifidobacterium longum* which were also found as reproducible across the datasets from our meta-analysis, also ranked among the top 15 discriminatory features. Altogether, we obtained substantial amount of overlap between species-level meta-analysis (−0.2 > effect size > 0.2; FDR < 0.1) and LODO machine learning experiments which includes a total of 12 species. Together this species set comprise the biomarkers of obese gut microbiome (Figures 3B and 5C and Table S4.1). Further, we performed LODO analysis exclusively with these biomarkers to observe if they can attain similar accuracy value as obtained using all features to get stronger evidence in support of their obesity-specificity. Interestingly, we noticed that these features were not only sufficiently discriminatory but also showed an improved accuracy in majority of the datasets (China, Denmark, Ireland and Japan) (Fig 4A). Among the other datasets, for GBR (AUC = 0.67) and Kazakhstan (AUC = 0.67), the biomarkers sufficiently discriminated between the classes with the only exception for Sweden (AUC = 0.64). Altogether, LODO-generated model using relative abundance of the few selected biomarkers successfully attained similar median accuracy (AUC = 0.67) as obtained by using all features (AUC = 0.68) (Figures 4A and 4C). Thus, substantial evidence was found for the species set to be called as robust biomarker associated with obesity associated.

Further, for pathway validation, we again selected 32 top-ranking species from each LODO run and averaged their ranking and compared to our pathway-level meta-analysis result. Expectedly, majority of the signature pathways with highest effect sizes were also reflected in the list of top 32 most discriminative pathway features (Table S6). However, unlike taxa, we did not find a consistency between the average feature ranking and their effect sizes.

### Reproducible functional shifts are driven by both dataset-specific and reproducible contributor species combinations

We observed a replicable signature pattern at both taxa and function levels by meta-analysis (Figures 3B and 3D). These patterns were obtained independently through comparative analyses of the obese and lean control groups. However, to understand contributor of the functional shift associated with obesity, these two feature levels remain to be precisely mapped which is crucial to get a glimpse of the biology of obesity and get an overview of the therapeutic intervention needed to revert the obesity associated dysbiosis. We attempted to establish this link with the help of FishTaco, a state-of-the-art computational framework that systematically integrates the taxonomy with the functional shifts (Manor and Borenstein, 2017). FishTaco exploits the gene or pathway copy number (genomic content) of each taxon and the feature abundance information to identify the “Direct”- and “Indirect”- contributors of a functional shift. Control-enriched taxa encoding a control enriched function are the “Direct contributors” of that function. In contrast, obese-enriched taxa which lack that particular function (or present at relatively low copy number) are termed as “Indirect contributors” of that function. (Manor and Borenstein, 2017). FishTaco was run on each independent dataset, and the contributors of each reproducible function obtained from pathway level meta-analysis were identified. We curated the top 5 direct and indirect contributors (Table S7.1) of all control- and obese-enriched reproducible functions for every dataset, and further analyses were performed.

For the analysis, we considered the presence of a contributor at least in 4 out of 7 datasets as the reproducibility criteria. Using this, we identified 10 such reproducible contributors driving various pathway shifts and visualized the pathways they contribute to (Figure 6A and Table S7.2). Strikingly, among the 10 reproducible contributors obtained in FishTaco analysis, all except *Prevotella copri* and *Eubacterium rectale* are direct contributors of control enriched pathways and drive most of the functions, which reiterate reduction of control-enriched taxa being the primary cause of the functional shifts. This finding is in perfect alignment with our observation of reduced diversity (Figure 2A) in obesity.

**Figure 6.**
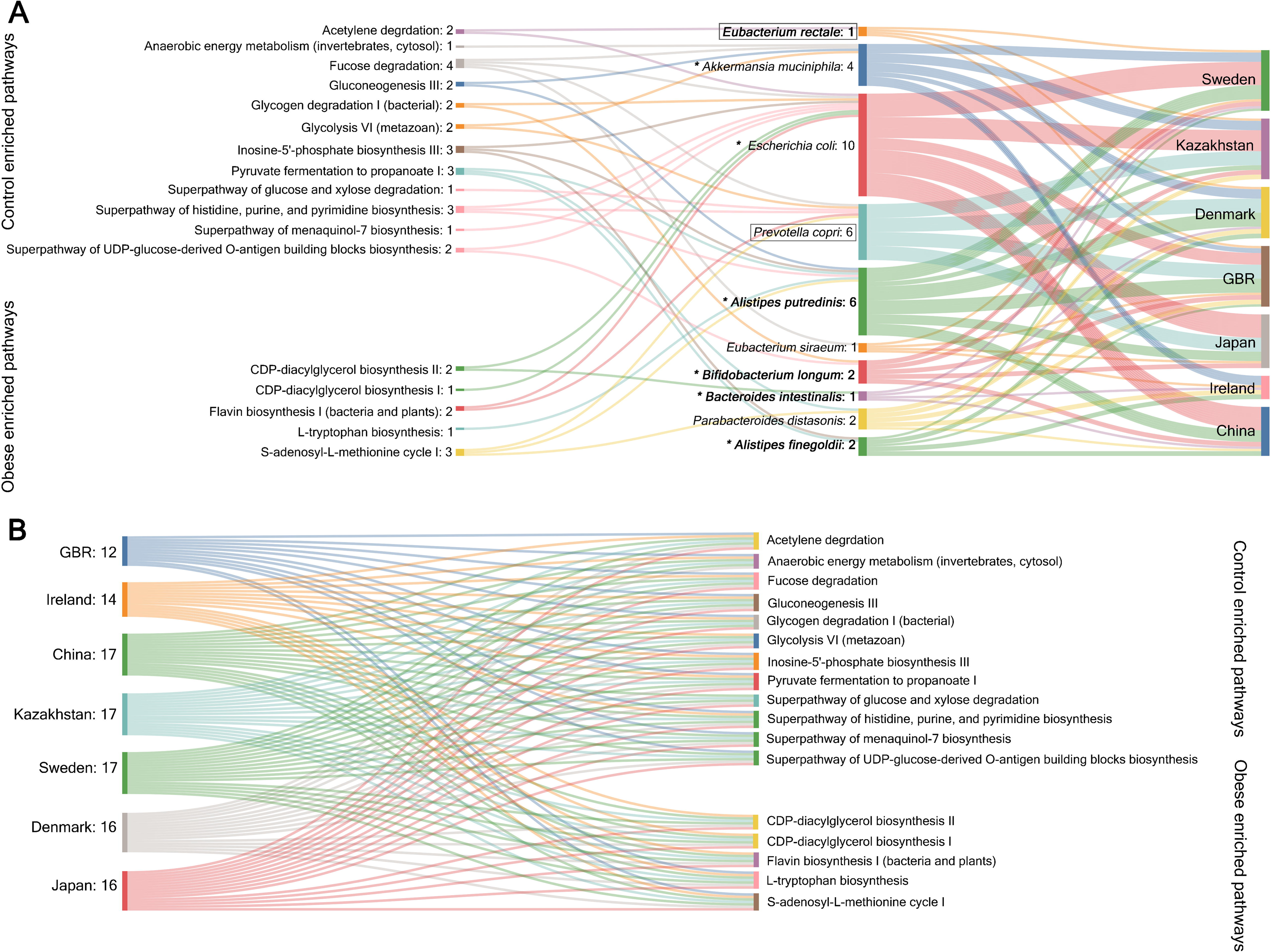
Reproducible contributors of the obesity-linked signature pathway shifts and predictive modulation of signature pathways by the minimal replicable contributor set. (A) Sankey diagram representing reproducible contributors (4 out of 7 datasets) and their contributed pathways. Values mentioned in each column represents the number of connections. Species marked in bold are the ones identified in both meta-analyses and MetaML and obtained as reproducible contributor through FishTaco. ‘*’ marked are the ones which are predicted as the minimal set of species contributing to the maximum number of reproducible pathways. (B) Sankey diagram shows probable control- and obese enriched signature pathways that can be predictively perturbed by minimal reproducible contributor set in different datasets. Values mentioned in each ‘dataset’ column represents the number of pathways the minimal set of species contributes.

We observed that majority of the species obtained in both meta-analyses and machine learning were also identified as reproducible contributors of at least one of signature pathways observed in pathway meta-analyses (Fig 6A). We denote this set of species (*Eubacterium rectale*, *Alistipes putredinis*, *Bifidobacterium longum*, *Bacteroides intestinalis*, and *Alistipes finegoldii* as biomarkers) (Figures 3B, 5C and 6A; Table S7.2). The rest of the contributors that were absent in meta-analyses and machine learning species, together with dataset-specific contributors, collectively drive many pathway shifts (Table S7.1). Among such reproducible contributors *Escherichia coli* is important being the driver of highest number of pathways (Figure 6A). Strikingly, some biomarkers (e.g., *Intestinimonas buyriciproducens*, *Alistipes indistinctus*) with high effect sizes (in meta-analysis) and low average ranks (machine-learning) were not found as reproducible contributors of signature pathway shifts. Thus, we suggest, in obesity, it is not enough to independently identify the reproducibly altered taxa through conventional analyses as many of them even do not reproducibly contribute to the functional shifts, while cumulative contribution of many insignificantly altered taxa might play remarkable roles in driving those shifts.

## Discussion

Obesity has been associated with multiple aetiologies including gut microbiome as one of the important links. Although many previous studies have specifically tried to demystify this link, there are several unresolved questions related to reproducibility in diversity, F/B ratio, taxonomic and functional features etc. and their association to obesity. In this study, we extensively assessed and evaluated several of these issues by analysing large-scale gut microbial metagenomic sequence data from 7 different geographical locations. Integration of samples from wide sources helps to take in to account the heterogeneities due to variation in geography, host genetics, ethnicity, age, sex, dietary pattern, physical activity, environment, birth mode, etc. Thus this strategy helps to identify obese-associated robust signature superseding the bias or noise introduced from batch effects and other confounders (Ma et al., 2014; Sung et al., 2013). From our study, we report a consistent reduction in alpha diversity in obese gut microbiome which is also reinforced by a previous 16s sequence-based meta-analysis (Figure 2A) (Sze and Schloss, 2016). This observation is quite relevant in the context of obesity as it has been rightly concluded that enriched gut microbial diversity is important for resilience and functional performance of intestinal gut microbial community (Clarke et al., 2014). Also, loss of diversity in the context of obesity has been compared to the ecosystem impoverishment in the macro ecology like a fertilizer runoff (Turnbaugh et al., 2008). In line with the species level diversity reduction, we also observed reduced gene family richness in obese individuals (Figure 2A). This could partly explain how a loss in microbial community diversity imparts a functional dysbiosis. Indeed, a previous report suggests effective resource utilisation by diverse gut microbial community, which in turn promotes a balanced and resilient microbial community (Lozupone et al., 2012). Further, high gut microbial diversity maintains functional redundancy in lean individuals, and reduced variation in obese may lead to disproportionate increase of selective adverse functions or loss of beneficial functions with respect to human host. Interestingly, Chatelier et al. reported a link between reduced gene richness and with higher BMI, insulin resistance, increased gut permeability, endotoxemia and inflammation (Le Chatelier et al., 2013). Further, in line with a previous pooled study (Sze et al., 2016), we observed higher gut microbial compositional dissimilarity in lean individuals. Additionally, similar to a previous 16S metagenomic study, we also conclude that F/B ratio is not a strong distinctive parameter for obese and the lean classification and thus cannot be used as a robust biomarker of obesity-associated gut microbial dysbiosis (Figure 2A) (Magne et al., 2020; Sze and Schloss, 2016).

We also established a panel of robust gut microbial species signatures by using random-effect meta-analysis which were further independently validated by a random forest-based machine learning classifier. Common species obtained from these two independent methods forms the basis to identify the biomarkers (Figures 3B and 5C and Table S4). The identified biomarkers are sufficiently able to reach near maximal prediction accuracy when validated on datasets independent from training the model (LODO) (Figures 4A and 4C). We expect further inclusion of more datasets with larger sample size would be useful for achieving even better prediction performance. It is worthwhile to mention that our LODO machine learning experiments captured the underlying signal more efficiently than a previous study performing CV with genus level abundance data (Sze and Schloss, 2016). This suggests that the obesity-associated biomarkers deciphered from species level data is more accurate, insightful and reliable. However, most of the biomarkers are control enriched and has been already established to have beneficial role against metabolic syndrome or negatively associated with high BMI. Among the biomarkers, *Intestinimonas butyriciproducens*, *Alistipes indistinctus, Alistipes putredinis, Alistipes shahii, Alistipes finegoldii*, are short chain fatty acids (acetic, isobutyric, isovaleric, propionic acid) producers. These bacteria through production of SCFAs (like acetate, butyrate, propanoate) promotes energy expenditure, insulin sensitivity, satiety hormone production and appetite regulation, reduces systemic low-grade inflammation and thus contribute to improved metabolic health (Canfora et al., 2015). Additionally, both *Bifidobacterium longum and Akkermansia muciniphila* obtained in our meta-analyses were reported to have beneficial role by promoting gut barrier function and preventing metabolic endotoxemia due to “leaky” gut (Cani et al., 2007).

However, scope to decode the mechanism of obese host-microbe association from species signatures is limited. Thus, from the metagenomically reconstructed pathways, we statistically analysed and revealed altered pathway signatures of obese gut microbiome. The control-enriched pathways mainly belong to purine and pyrimidine biosynthetic pathways fermentation pathways, carbohydrate metabolism pathways, nucleotide sugar biosynthesis pathways and menaquinol biosynthetic pathways (Figure 3D and Table S4). Our results suggest an enrichment of fermentation pathways which are possibly responsible for producing SCFAs like acetate, propionate and butyrate, as reported earlier (Caspi et al., 2014). On the contrary, biosynthesis pathway of flavin which is a cofactor of flavin containing monooxygenase 3 (FMO3) enzyme shows obese enrichment with highest effect size. Gut microbiota driven trimethylamine/trimethylamine-N-oxide/flavin-containing monooxygenase 3 (TMA/FMO3/TMAO) pathway is often associated to hyperglycemia, hyperlipidemia, atherosclerosis and obesity development (Schugar et al., 2017). Moreover, tryptophan amino acid biosynthesis may be linked to appetite and food intake stimulation through ghrelin and neuropeptide-Y release.

In this study, for the first time, microbiome and functions have been connected to identify drivers of obesity-associated functional shifts. From our analyses, we found, that the contributors are both dataset-specific as well as reproducible across the datasets (Figure 6A and Table S7.1 and S7.2). Majority of the reproducible contributors were also found to be obese-associated biomarkers as deciphered from meta-analysis and machine learning experiments (Figures 3B, 5C, and 6A; TableS4). Additionally, we noticed, other reproducible contributors together with dataset-specific contributors despite their insignificant alteration in abundance drives the obesity-associated gut microbial pathways. Thus, we infer from our study that although same set of functional alterations exist across datasets, these are not necessarily contributed by same set of taxa. Thus, mere identification of differentially abundant species and looking into its functional potential is not enough for understanding the biology and strategizing therapy rather the complex interconnectivity with the gut microbial ecosystem should be carefully considered. Additionally, we aimed at predicting a minimal set of species that can contribute to maximum number of reproducible pathway shifts in different datasets. For that, we suggest the overlap of the reproducible contributors and the reproducible taxa obtained from meta-analysis together with *Escherichia coli* as it is the most critical reproducible contributor. Together this group of 6 species are predicted to collectively drive all 17 reproducible functions (12 control enriched and 5 obese enriched pathways) obtained in meta-analysis (Figure 6B, Table S7.3). It is to be noted that being the indirect contributor of the obese enriched pathways, these group if enriched in the obese might ultimately cause depletion of these pathways. Therefore, in presence of other beneficial species these 6 contributors might play extremely important role in reverting the obese associated dysbiosis.

Our study has some limitations. Although we minimized gut microbial variations arising due to differences in sequencing platform and comorbidity factors, several other factors limit power of our study like host genetics, diet etc. that we were unable to access due to unavailability of the relevant metadata. Also, our random forest-based classifier has limited power in deciphering functional pattern and thus deep learning-based classification may be used for obtaining better discriminatory pattern. Also, in the context of our study, intestinal mucosa-associated microbiome and virome data which are currently unavailable can further enrich the understanding of microbiome association to obesity. Further including microbial metabolites in this study could add additional dimensions towards connecting gut microbiota to obesity.

## Acknowledgement

This study is supported by Start-up Research Grant (SRG/2019/000221) from Science & Engineering Research Board (SERB), Government of India to D.D. D.C also extends his gratitude to Council of Scientific & Industrial Research (CSIR), New Delhi, India, for their financial supports. Both the authors acknowledge Mr. Ashis Roy and Mr. Monojit Kamilya for their valuable inputs in with the computational analyses.

## Author Contributions

D.C. collected the data, carried out the analyses according to the adopted pipeline and wrote the paper. D.D. framed the research question, analysed the data, supervised the work and wrote the manuscript. Both the authors approved the final version of the paper.

## Figure Legends

**Figure S1. Flow diagram of the dataset search and sample inclusion strategy used in our study**

**Figure S2, related to** **Figures 2. Differences in taxonomic and functional structures of gut microbial community between obese and controls**

(A-E) Notched boxplots reporting (A) Shannon- and (B) Simpson-species diversity index, (C) gene family richness, (D) Firmicutes/Bacteroidetes ratio, and (E) beta-diversity index calculated after rarefaction (See Methods) in each dataset. P values were calculated by two-tailed Wilcoxon rank-sum tests.

**Figure S3, related to** **Figures 2A. Multivariate analysis of species and functional diversity indices**

(A-C) Multivariate analysis of (A) Shannon- and (B) Simpson-species diversity index, (C) gene family richness using crude and age-, sex-adjusted coefficients obtained from linear models using MaAsLin2 for each dataset.

(D-E) Meta-analysis of crude and age, sex adjusted coefficients obtained from multivariate analysis for each dataset done with the species diversity indices (D) and gene family richness (E) using a random effects model. Red lines represent the 95% confidence interval for the random effects model estimate.

**Figure S4, related to** **Figures 3A. Linear Discriminant Analysis Effect Size (LEfSe) of microbial taxa between obese and controls**

LEfSe analysis of species with LDA score cut off > 2.0 and < -2.0 (p-value <0.05) was considered as significantly differentially abundant between obese and control groups. Species were ranked by their respective LDA score. LEfSe plots of taxonomic biomarkers were generated on Galaxy computational tool v.1.0. (https://huttenhower.sph.harvard.edu/galaxy/).

**Figure S5, related to** **Figures 3C. Linear Discriminant Analysis Effect Size (LEfSe) of microbial functions between obese and controls**

LEfSe analysis of pathways with LDA score cut off > 2.0 and < -2.0 (p-value <0.05) was considered as significantly differentially abundant between obese and control groups. Pathways were ranked by their respective LDA score. LEfSe plots of functional biomarkers were generated on Galaxy computational tool v.1.0. (https://huttenhower.sph.harvard.edu/galaxy/).

**Figure S6, related to** **Figures 3B** **and 3D. Multivariate analysis of signature features obtained from meta-analyses to access the effect of potential confounders.**

(A) Multivariate analysis of gut microbial signature species associated with obesity health status using crude and age, sex adjusted coefficients obtained from linear models using MaAsLin2.

(B) Random effects meta-analysis on crude and age, sex adjusted coefficients obtained from multivariate analysis of microbial species signatures linked to obesity using MaAsLin2. Red lines represent the 95% confidence interval for the random effects model estimate.

(C) Multivariate analysis of signature gut microbial pathways linked to obese phenotype (See Figure 3D) using crude and age, sex adjusted coefficients obtained from linear models using MaAsLin2.

(B) Random effects meta-analysis on crude and age, sex adjusted coefficients obtained from multivariate analysis of microbial pathway signatures associated with obesity using MaAsLin2. Red lines represent the 95% confidence interval for the random effects model estimate.

**Figure S7, related to** **Figures 4A. LODO prediction performance with increasing number of training datasets**

(A-G) Prediction accuracy computed with species relative abundance data with subsequent increase in the number of training datasets in LODO experiments for each validation datasets. Each point shows median AUC values obtained from corresponding number of training datasets trained in every possible combination.

(H) Matrix shows median AUC values with increasing number of training datasets for corresponding validation dataset present in the row.

## Supplemental table legends

**Table S1, related to** **Figure 1**. Description of datasets included in this study

**Table S2, related to Figure S2:** P-values obtained from univariate non-parametric analysis of Alpha-, beta- diversity and Firmicutes/Bacteroidetes ratio between obese and control groups.

**Table S3, related to** **figure 2**. Results of meta-analyses on alpha-, beta-diversity and Firmicutes/Bacteroidetes ratio.

**Table S4, related to** **Figure 3**. Species and functional pathway-level meta-analysis results.

Table S4.1, related to Figure 3B. MetaPhlAn3 species-level meta-analysis result.

Table S4.2, related to Figure 3D. HUMAnN3 pathway-level meta-analysis results (FDR<0.05)

Table S4.3, related to Figure 3D. Signature pathways with highest effect sizes (Effect Size cut off ±0.35) obtained from meta-analysis

**Table S5, related to** **Figures 4E** **and 4F.** P-values (calculated with Wilcoxon rank-sum test) between the median AUC values obtained using original and shuffled class labels.

**Table S6, related to** **Figure 5B**. Pathways ranked within top 32 features in at least one dataset are reported along with their median and final ranks. Pathways marked in bold and colour-filled cells were identified from pathway-level meta-analysis (See Figure 3D).

**Table S7, related to** **Figures 6**. Results of FishTaco analyses.

Table S7.1, related to Figure 6A. Dataset-specific top 5 direct- and indirect- contributors of the signature pathways associated with obese health phenotype. Reproducible contributors (contributing a pathway in at least 4 out of 7 datasets) are shown in bold.

Table S7.2, related to Figure 6A. Reproducible contributors of obesity-linked signature pathways (curated from Table S7.1).

Table S7.3, related to Figure 6B. Probable obesity-linked signature pathways that may be partially perturbed by suggested minimal species set *(Alistipes finegoldii*, *Alistipes putredinis*, *Bifidobacterium longum*, *Bacteroides intestinalis*, *Akkermansia muciniphila*, *Escherichia coli*) in different datasets.

