## Supplemental Figure 1 for "Meta-analysis reveals obesity associated gut microbial alteration patterns and reproducible contributors of functional shift"

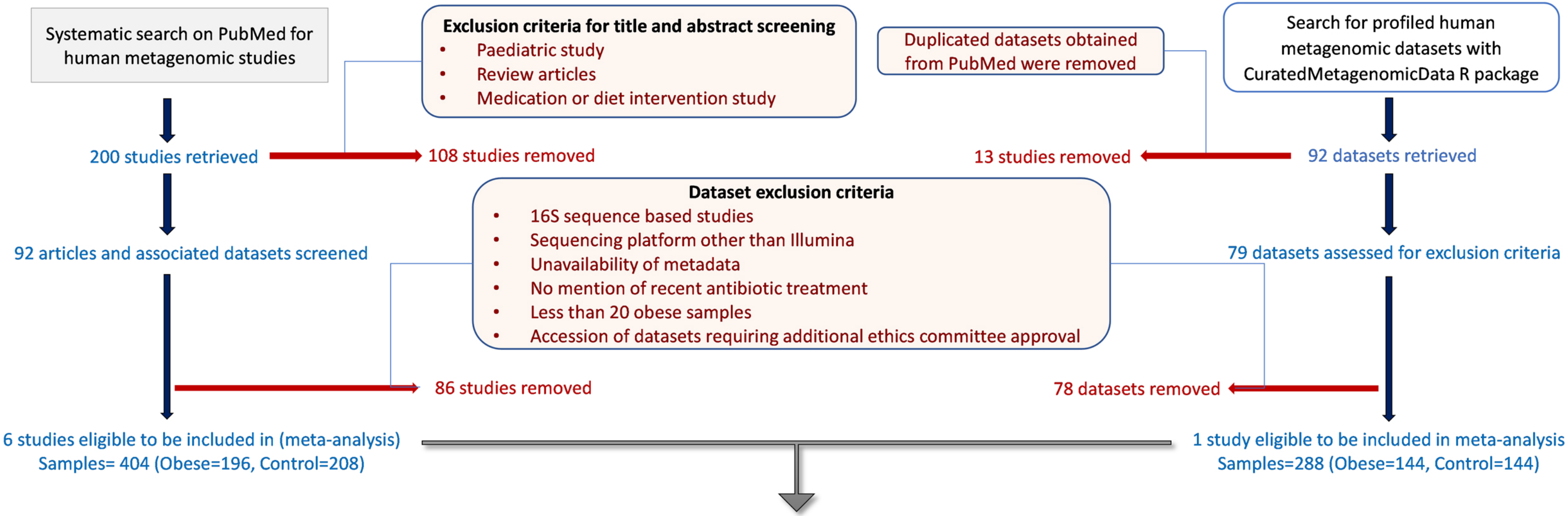


**Dataset used in meta-analysis**

| # | Datasets | ProjectID | # of controls | # of obese | Total samples |
|---|---|---|---|---|---|
| 1 | China | PRJEB21528 | 45 | 38 | 83 |
| 2 | Denmark | PRJEB2054 | 37 | 31 | 68 |
| 3 | Great Britain (GBR) | PRJEB39223 | 144 | 144 | 288 |
| 4 | Ireland | PRJEB37017 | 39 | 39 | 78 |
| 5 | Japan | PRJDB4176 | 46 | 45 | 91 |
| 6 | Kazakhstan | PRJEB17632 | 25 | 23 | 48 |
| 7 | Sweden | PRJEB1786 | 16 | 20 | 36 |
| Total 692 | | | | | |
