## Supplementary figures and images for "Meta-analysis reveals obesity associated gut microbial alteration patterns and reproducible contributors of functional shift"

### Supplemental Figure 2

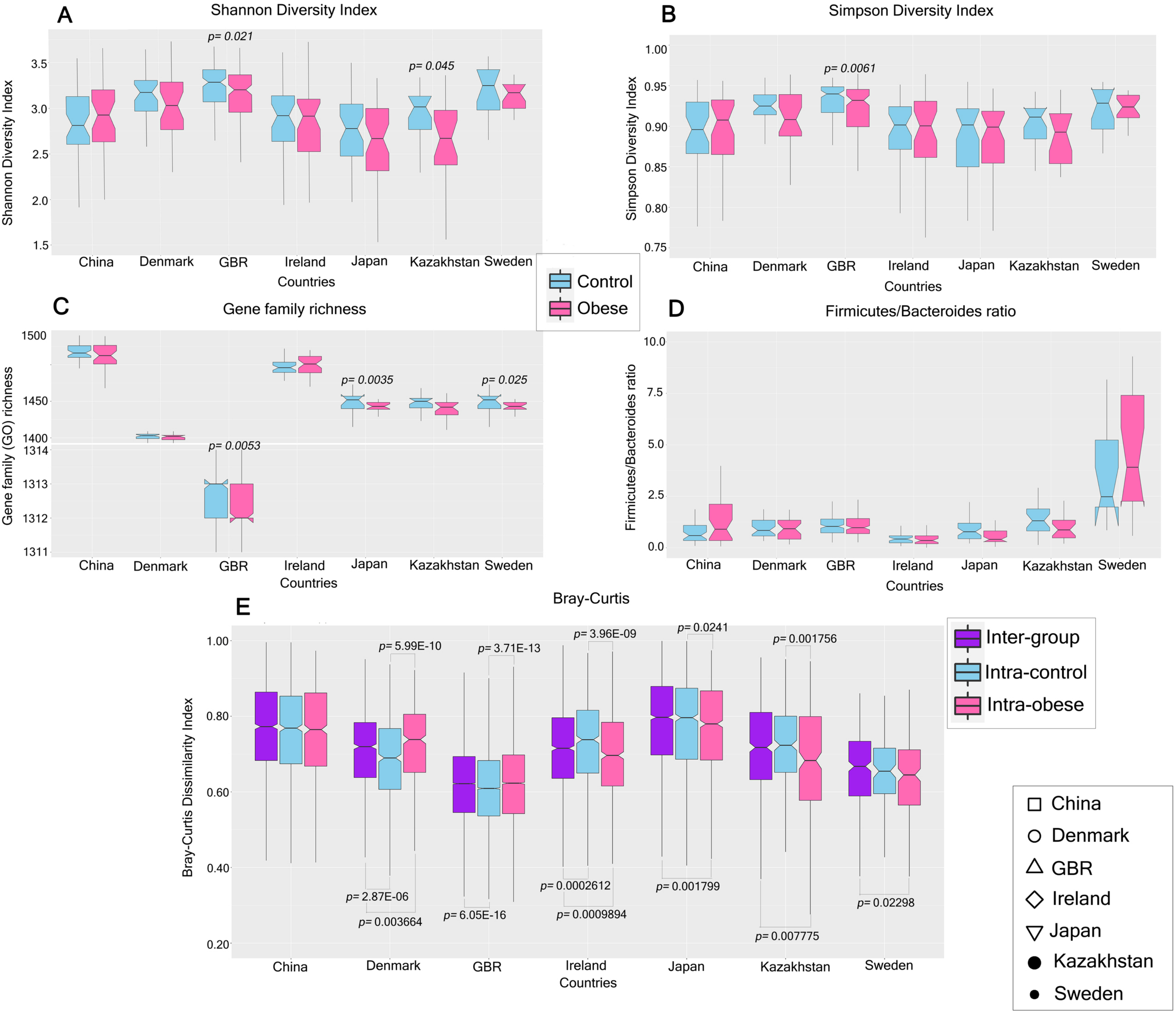

### Supplemental Figure 3

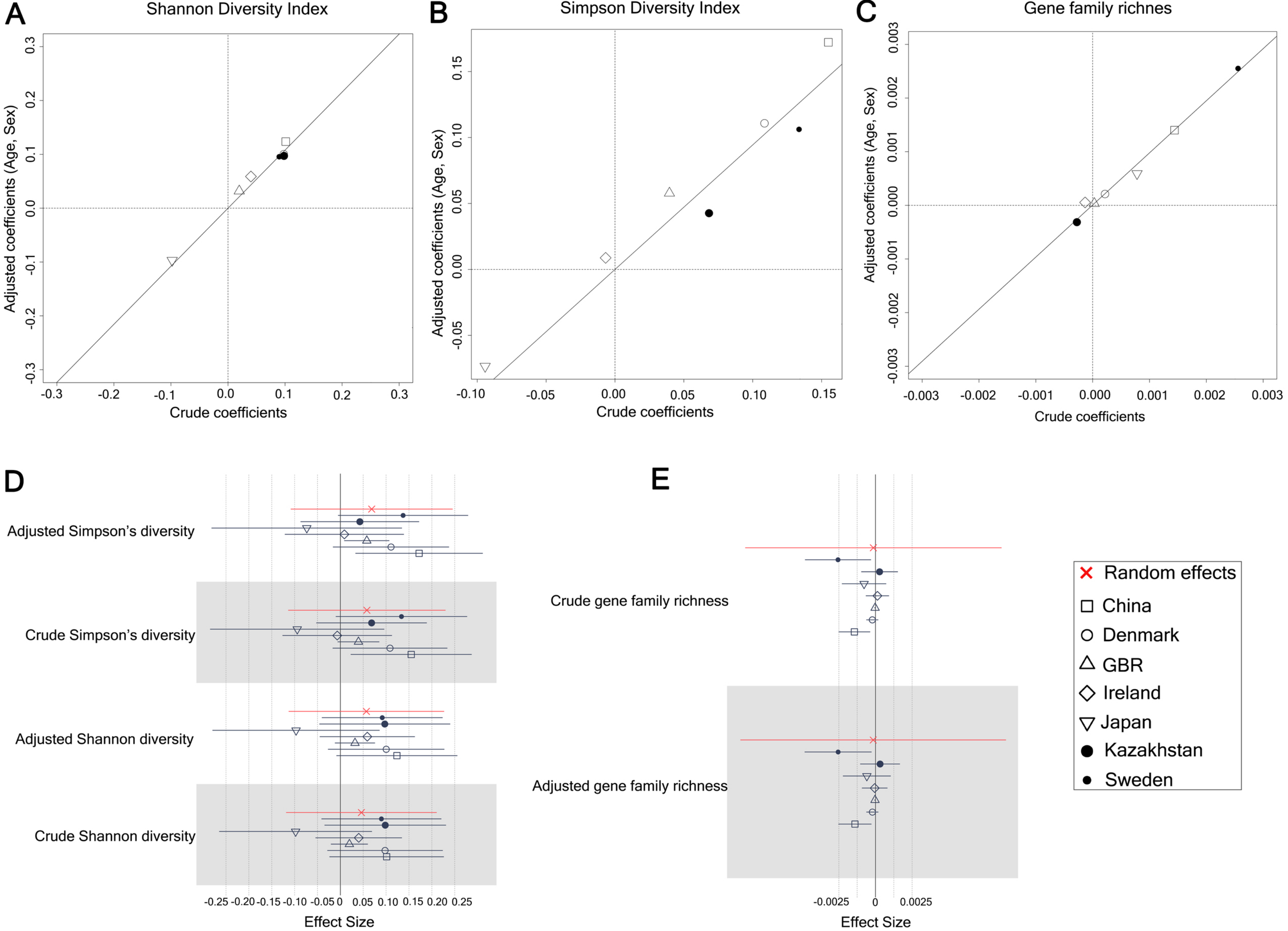

### Supplemental Figure 4

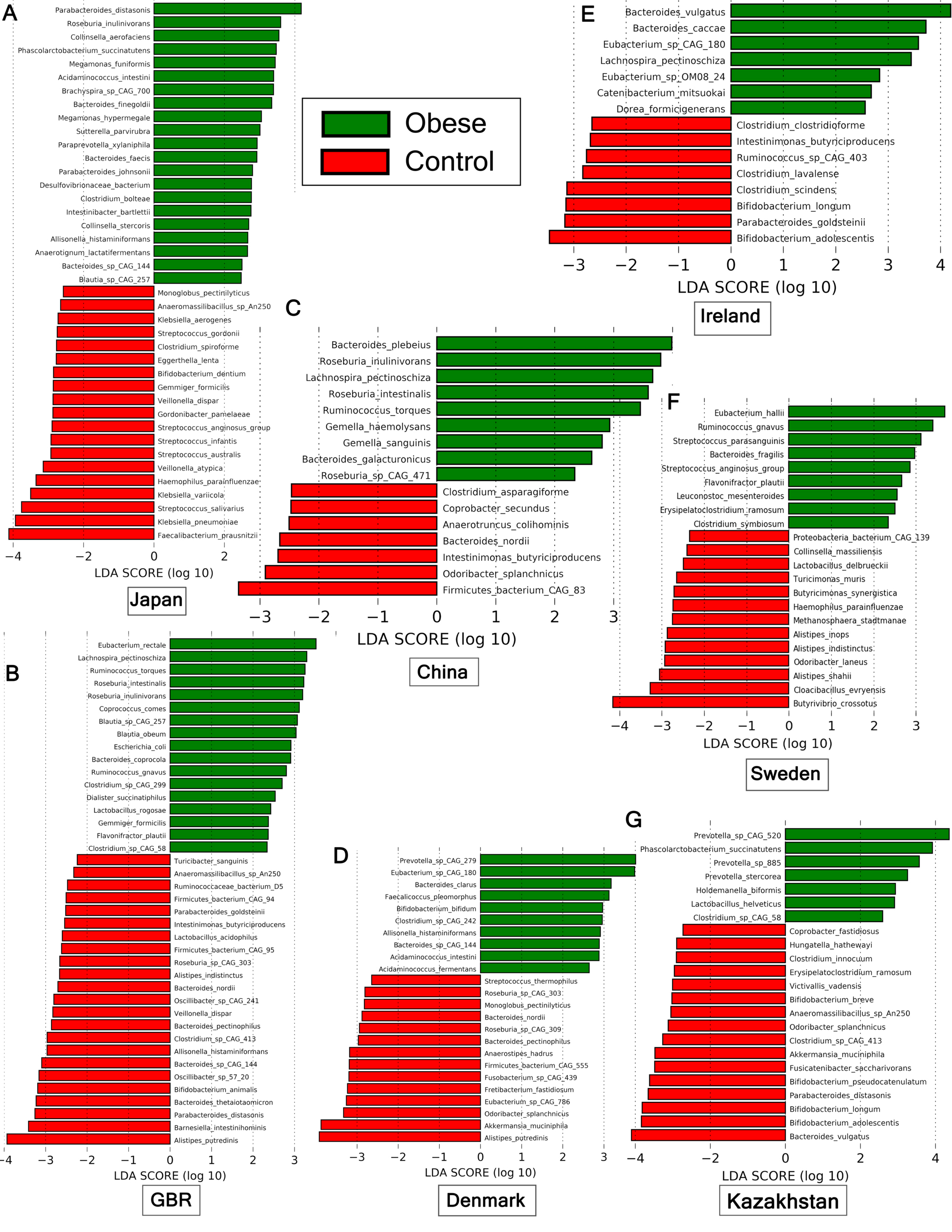

### Supplemental Figure 5

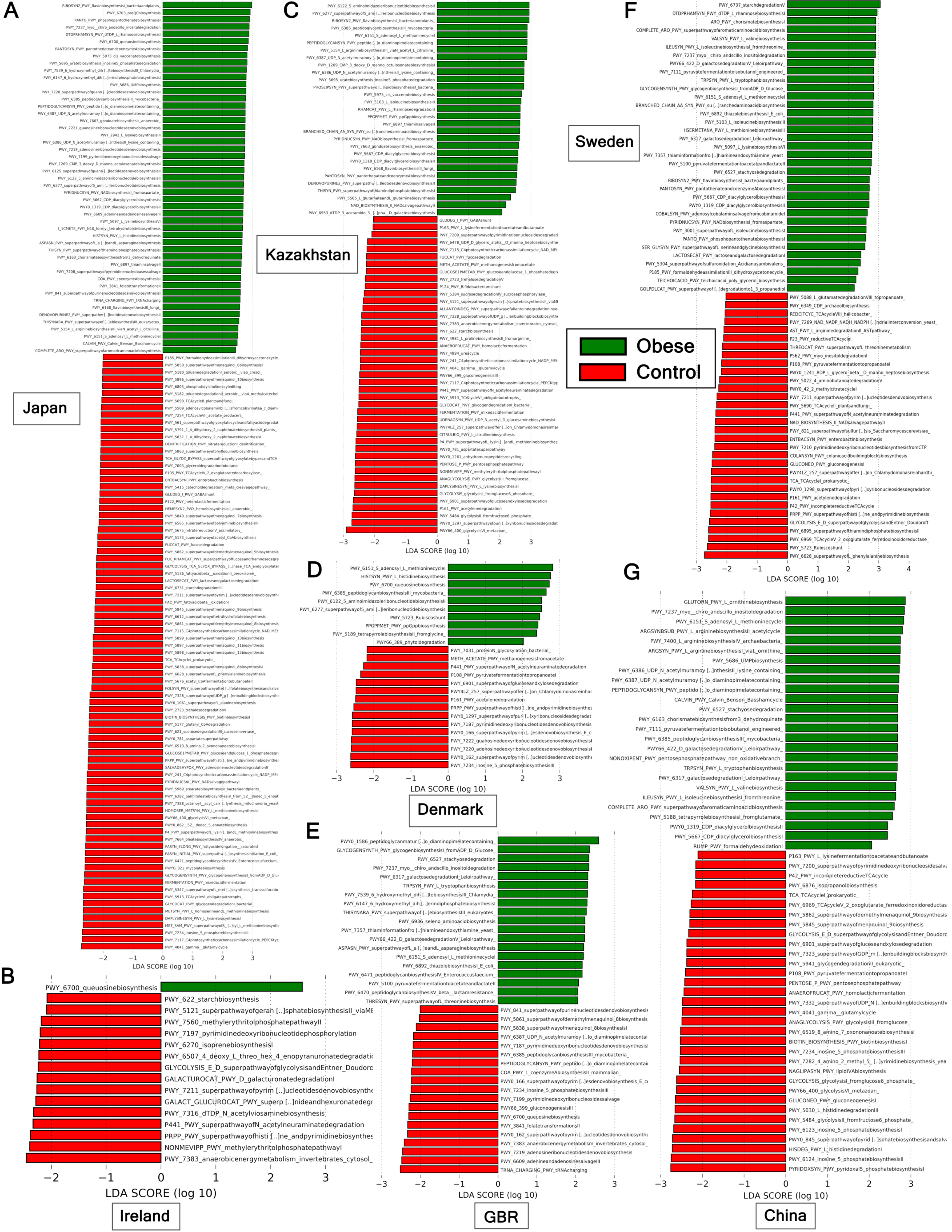

### Supplemental Figure 6

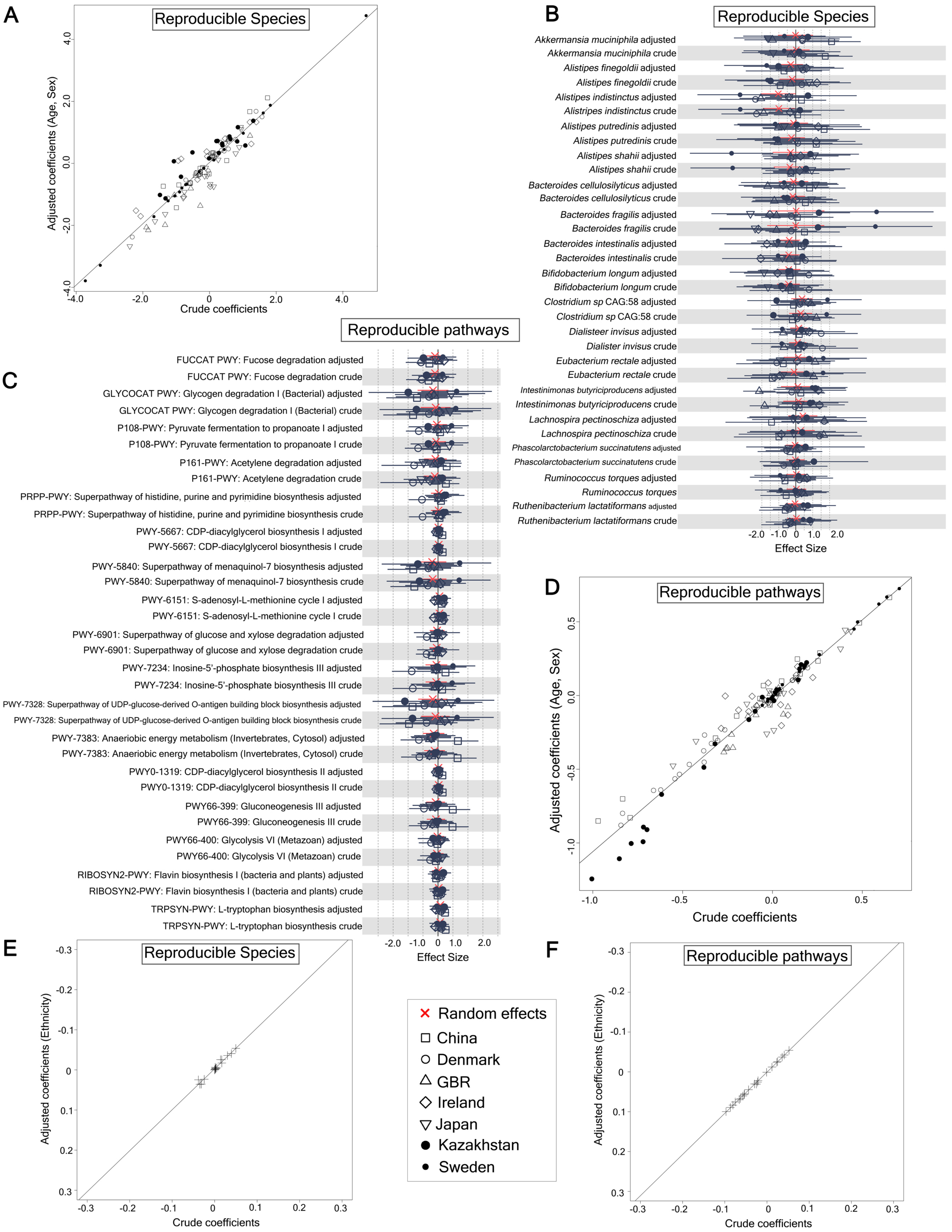

### Supplemental Figure 7

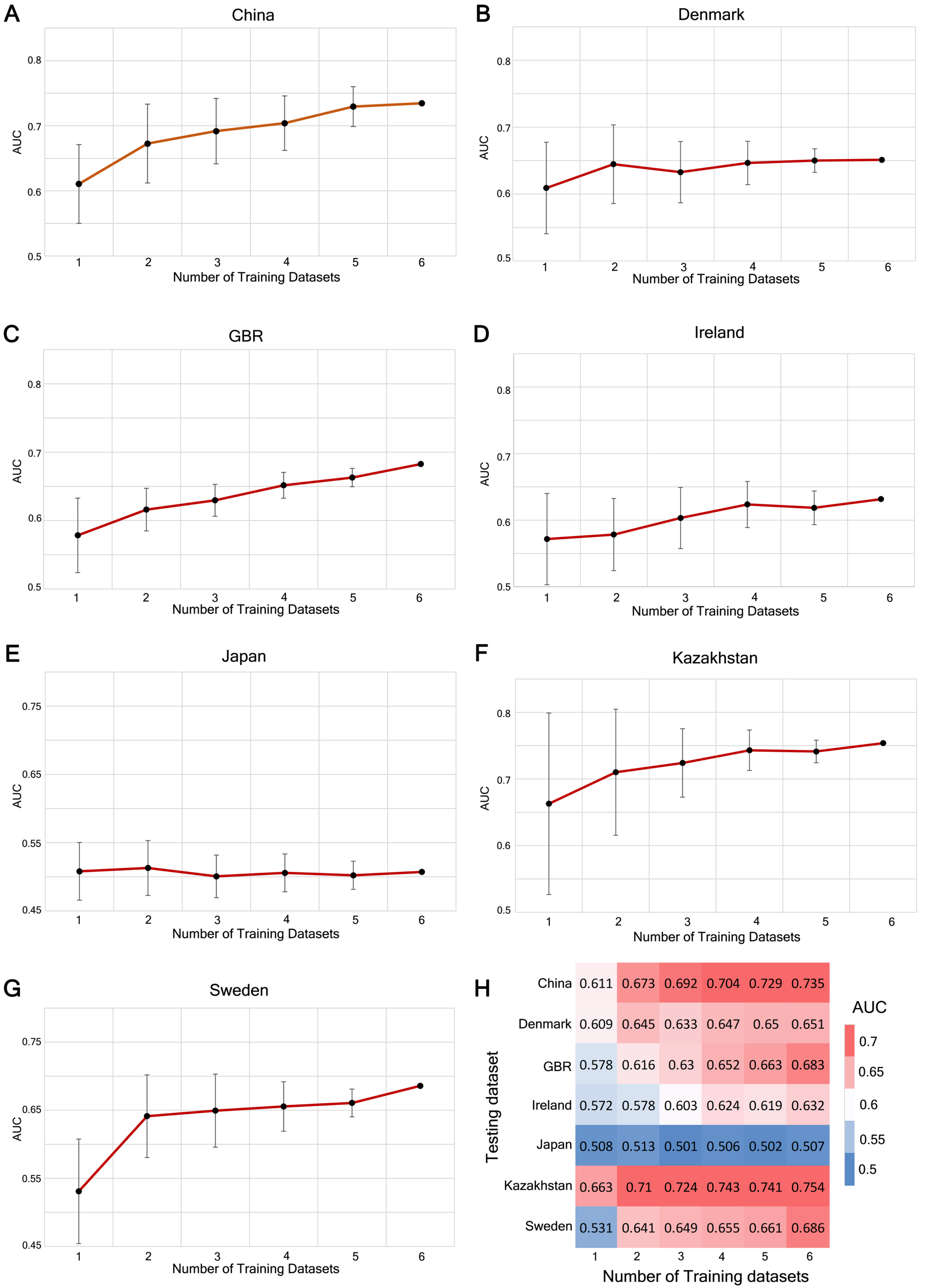
